# A pangenome and pantranscriptome of hexaploid oat

**DOI:** 10.1101/2024.10.23.619697

**Authors:** Raz Avni, Nadia Kamal, Lidija Bitz, Eric N. Jellen, Wubishet A. Bekele, Tefera T. Angessa, Petri Auvinen, Oliver Bitz, Brian Boyle, Francisco J. Canales, Brett Chapman, Harmeet Singh Chawla, Yutang Chen, Dario Copetti, Viet Dang, Steven R. Eichten, Kathy Esvelt Klos, Amit Fenn, Anne Fiebig, Yong-Bi Fu, Heidrun Gundlach, Rajeev Gupta, Georg Haberer, Tianhua He, Matthias H. Herrmann, Axel Himmelbach, Catherine J. Howarth, Haifei Hu, Julio Isidro y Sánchez, Asuka Itaya, Jean-Luc Jannink, Yong Jia, Rajvinder Kaur, Manuela Knauft, Tim Langdon, Thomas Lux, Sofia Marmon, Vanda Marosi, Klaus F.X. Mayer, Steve Michel, Raja Sekhar Nandety, Kirby T. Nilsen, Edyta Paczos-Grzęda, Asher Pasha, Elena Prats, Nicholas J. Provart, Adriana Ravagnani, Robert W. Reid, Jessica A. Schlueter, Alan H. Schulman, Taner Z. Sen, Jaswinder Singh, Mehtab-Singh, Nick Sirijovski, Nils Stein, Bruno Studer, Sirja Viitala, Shauna Vronces, Sean Walkowiak, Penghao Wang, Amanda J. Waters, Charlene P. Wight, Weikai Yan, Eric Yao, Xiao-Qi Zhang, Gaofeng Zhou, Zhou Zhou, Nicholas A. Tinker, Jason D. Fiedler, Chengdao Li, Peter J. Maughan, Manuel Spannagl, Martin Mascher

## Abstract

Oat grain is a traditional human food rich in dietary fiber that contributes to improved human health. Interest in the crop has surged in recent years owing to its use as the basis for plant-based milk analogs. Oat is an allohexaploid with a large, repeat-rich genome that was shaped by subgenome exchanges over evolutionary timescales. In contrast to many other cereal species, genomic research in oat is still at an early stage, and surveys of structural genome diversity and gene expression variability are scarce. Here, we present annotated chromosome-scale sequence assemblies of 33 wild and domesticated oats along with an atlas of gene expression across six tissues of different developmental stages in 23 accessions. We describe the interplay of gene expression diversity across subgenomes, accessions and tissues. Gene loss in the hexaploid is accompanied by compensatory up-regulation of the remaining homeologs, but this process is constrained by subgenome divergence. Chromosomal rearrangements have significantly impacted recent oat breeding. A large pericentric inversion associated with early flowering explains distorted segregation on chromosome 7D and a homeologous sequence exchange between chromosomes 2A and 2C in a semidwarf mutant has risen to prominence in Australian elite varieties. The oat pangeome will promote the adoption of genomic approaches to understanding the evolution and adaptation of domesticated oats and will accelerate their improvement.

---

Oat (*Avena sativa,* 2n = 6x = 42, A_s_A_s_C_s_C_s_D_s_D_s_ genome^1^) is the world’s seventh most widely grown cereal crop^2^. It is appreciated for its high content of dietary fiber, which has been shown to have substantial benefits for human health^3,4^, and as the raw material for many foods, including plant-based milk substitutes. In 2022/23, more than 25 million metric tons were produced worldwide. Genetically improved cultivars have the potential to make oat cultivation more productive and sustainable, but much of this potential remains unrealized. While wheat and barley researchers have had reference genome sequences of their species of interest for about a decade, oat has lagged behind, with the first oat reference sequences only published in the last few years^5–7^. The complexity of the oat genome is partly to blame for the slow progress. Oat is an allohexaploid species with the subgenomes A, C, and D, each between 3 and 4 Gb in size^5^. In contrast to bread wheat, which arose as a hexaploid only about 12,000 years ago^6,8^, oat’s conspecific wild progenitor *A. sterilis*^9,10^, a wild grass common in Western Asia and Mediterranean basin, has been a hexaploid for at least 500,000 years^11^. Inheritance in oat is disomic, i.e. chromosomes from different subgenomes (homeologs) do not generally recombine. Even so, the presence^6,11^ of three subgenomes in the same nucleus has afforded opportunities for rare homeologous exchanges to reshuffle the oat genome^5^. In the first analyses of a chromosome-scale oat genome sequence, Kamal et al.^5^ assigned chromosomes to the A, C and D subgenomes according to which diploid progenitor their pericentromeres were descended from. However, this was an incomplete approximation of oat genomic ancestry. For example, genes that could be traced back to C genomes now reside on chromosomes whose pericentromeres match those of A and D genome species^5^. Moreover, all but one intergenomic interchange occurred between the C and D subgenomes in the evolution of the tetraploid progenitor *A. insularis* (C_i_C_i_D_i_D_i_), which may well have existed for several million years prior to hexaploidization and the addition of genome A from *A. longiglumis*^5^. Now that contiguous genome sequences can be assembled even for complex, repeat-rich plant genomes^12^, the polyploid and mosaic ancestry of oat should no longer be seen as a challenge, but rather as an opportunity for pangenomic analyses. In this context, oat provides a model system for questions such as: How does genic presence-absence variation (PAV) affect gene expression in a polyploid? Does structural variation in an old polyploid, especially sequence exchange between subgenomes, impact breeding? To tackle these and other questions, we have studied gene expression diversity and structural variation in an annotated pangenome of cultivated oats and allied taxa.

## An annotated pangenome of hexaploid oat

We assembled and annotated the genomes of 33 diverse oat lines (**Supplementary Table 1**). Henceforth, we refer to these lines as the PanOat panel. This panel comprises: (i) commercially successful elite varieties from major oat-growing regions; (ii) plant genetic resources with interesting properties; (iii) two accessions of wild *A. sterilis*; (iv) the closest extant relatives of oat’s diploid and tetraploid progenitors, *A. longiglumis* and *A. insularis;* (v) Amagalon, a synthetic hexaploid; (vi) two distant diploid *Avena* species *A. eriantha* and *A. atlantica*. The PanOat lines cover the genetic diversity space of the crop, as represented in a principal component analysis (PCA) of 9,174 diverse wild and domesticated genebank accessions **(Fig. 1a)**. Genome sequences for six members of the PanOat panel were published previously^5,7,13^. The genomes of three lines, Gehl, AAC Nicolas and *A. byzantina* PI258586, were sequenced with Illumina short reads. The remaining 24 genome sequences were assembled from accurate long-reads generated on the PacBio HiFi platform (**Supplementary Table 1**). All contig-level assemblies were scaffolded with chromosome conformation capture sequencing (Hi-C) data. On average, 99.97% of the assembled sequences were assigned to precise chromosomal locations (**Supplementary Table 1**). We annotated genes on these assemblies using a multi-tiered approach^14^. To do so, we sequenced the transcriptomes of six different tissues and developmental stages in 23 PanOat lines (**Fig. 2a, Extended Data Fig. 1, Extended Data Fig. 2a,b, Supplementary Table 2**) using Illumina RNAseq. In addition, we sequenced pooled samples using PacBio Iso-Seq. For two additional lines (*A.sterilis* - TN4 and *A. byzantina* PI 258586) we used only RNAseq data. These data, as well as evidence from protein homology and *ab initio* predictions, were used to predict gene models, which were then projected onto the eight PanOat assemblies without native transcriptome data. All gene models were assigned to the high- and low-confidence categories^15^ based on their homology to genes in other plants and the presence of domains commonly found in transposable elements (TEs). For a hexaploid oat genome, we predicted between 107,847 and 136,836 genes (**Extended Data Fig. 3a**), of which 60.5% on average were expressed (**Supplementary Table 2**). We used Benchmarking Universal Single-Copy Orthologs (BUSCO^16^) to assess the completeness of our gene annotations. Out of the 4,896 single-copy near-universal orthologs in the Poales BUSCO dataset (poales_odb10), an average of 4,839 (98.8%) were complete in hexaploid oats. Specifically, 3,680 (75.2%) were triplicated, 666 (13.6%) were duplicated, and 166 (3.4%) occurred in a single copy (**Supplementary Table 3, Extended Data Fig. 3b**). Completeness was slightly lower in short-read compared to long-read assemblies.

**Figure 1:**
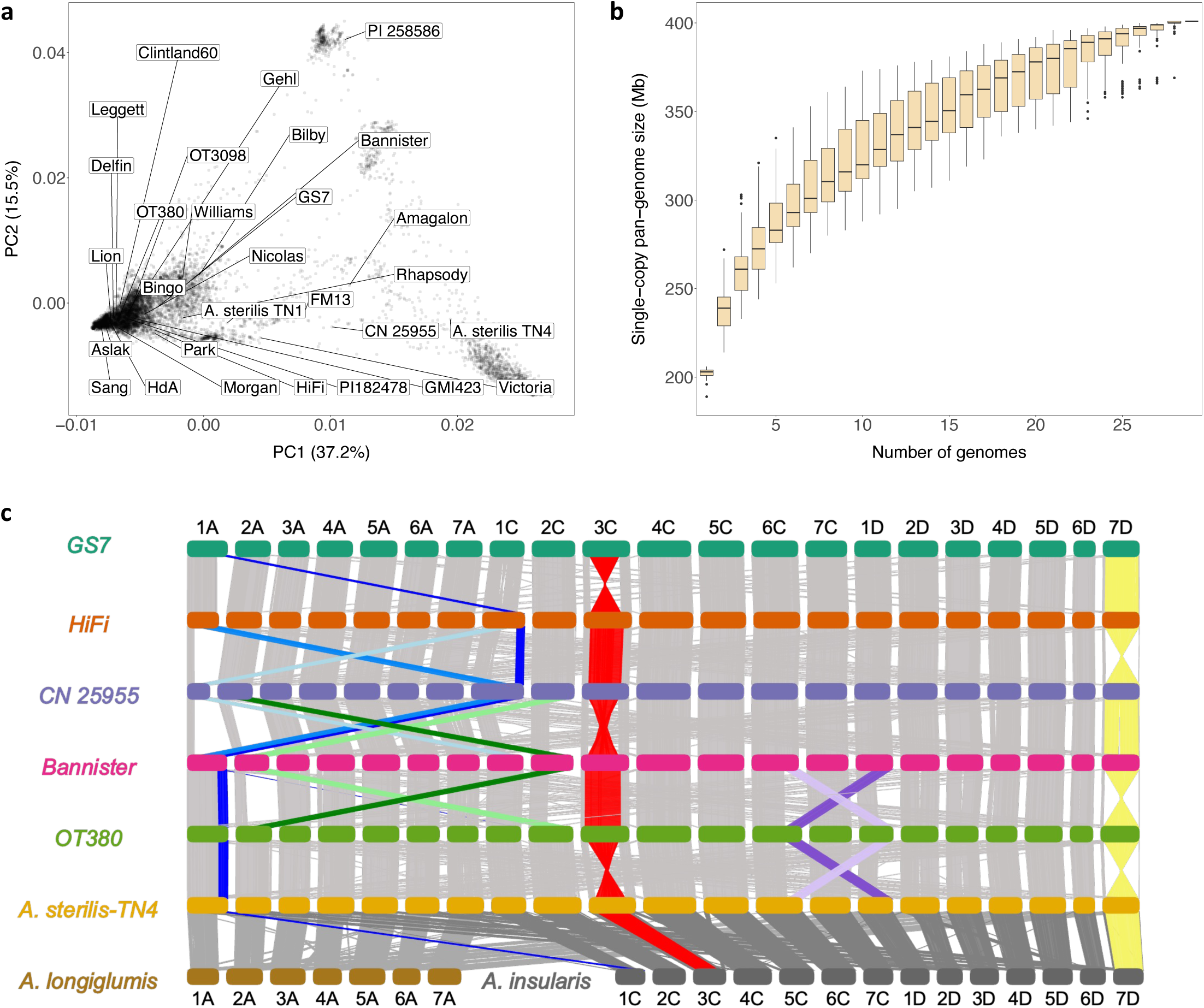
Oat pangenome composition and structural variation. **(a)** PCA plot of 9,174 hexaploid oats including A .sativa, A. byzantina and A. sterilis overlaid with the position of PanOat assemblies. **(b)** Hexaploid pangenome complexity estimated by single-copy k-mers. The curves trace the growth of non-redundant single-copy sequences as sample size increases. Error bars are derived from 100 ordered permutations each. (c) NGenomeSyn synteny plot between representative PanOat assemblies including the diploid and tetraploid genome donors (A. longiglumis – A genome and A. insularis – CD genomes) showing major SVs. The grey bars represent syntenic regions between all assemblies and the coloured bars mark the SVs in the different assemblies. *In panels a and c PI 258586 = A. byzantina PI258586, HdA = Hâtives des Alpes, Nicolas = AAC Nicolas, Morgan = AC Morgan, and CN 25955 = A. occidentalis CN 25955.

**Figure 2:**
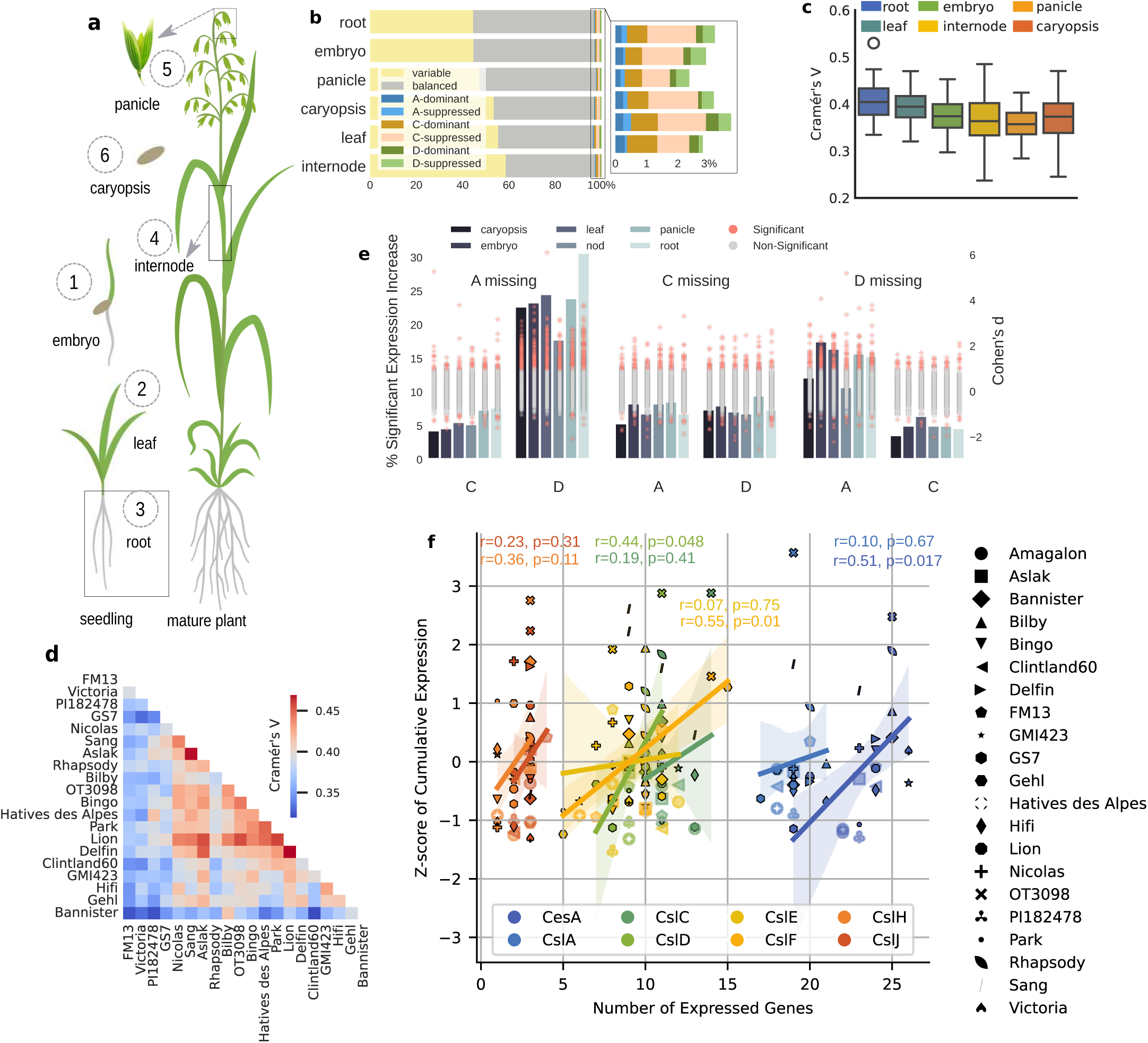
An oat pan-transcriptome. **(a)** Illustration of the harvested tissues for transcriptome sequencing. **(b)** Proportions of stable and variable genes across 20 Avena sativa lines in six different tissues. For stable genes, the expression categories are classified into A-, C-, and D-dominant and A-, C- and D-suppressed, and balanced. **(c)** Distance matrix based on expression level categories in 20 Avena sativa lines, using Cramér’s V, exemplified by root tissue. **(d)** Pairwise distribution of Cramér’s V values between different tissues, indicating the degree of similarity in gene expression patterns. **(e)** Dynamics of gene expression inthe comparison of complete triads to those lacking a member from one subgenome. This panel shows the percentage of cases with significant expression increases in each of the other two subgenomes when one homeolog is missing, along with the corresponding Cohen’s d values. **(f)** Gene expression and gene copy number variation of cellulose synthase and cellulose synthase like genes in the oat pangenome, illustrating the correlation between the gene expression and copy number variation.

## A catalogue of gene-based presence-absence variation

The gene content of a pangenome can be divided into core, shell, and cloud compartments, which consist of genes present in all, many, or a single line, respectively^17^. To study genic presence-absence variation (PAV), we constructed an orthologous framework from our gene annotations using Orthofinder^18^. To define the three genic categories — core, shell, and cloud — in the 30 members of the PanOat panel that are part of this framework (**Supplementary Table 3**), we used the following thresholds: core genes are present in 30 genomes; shell genes in 2-29 and cloud genes in a single genome. Our orthologous framework had 102,076 Hierarchical Orthologous Groups (HOGs). The core genome comprised 12,671 HOGs (943,786 genes) that contained at least one orthologous gene from all 30 lines. In total, we found 32.7% of the genes in the core-, 66.2% in the shell-, and 1.1% in the cloud-genome (**Extended Data Fig. 3c,d, Supplementary Table 3**). By definition, cloud and shell genes are not present in one or more genomes; we observed PAVs with varying contributions across different genomes (**Extended Data Fig. 3c,d**). The core genome was enriched for genes involved in essential physiological processes such as flower formation, nutrient uptake and cell wall organisation (**Supplementary Table 4)**. In contrast, the shell genome was enriched for genes related to defense mechanisms and activities as well as seed storage processes. It included genes encoding for various transcription factor families such as MYB, WRKY, NAC, AP2/ERF, and MADS/AGL. The cloud genome was notably enriched for phosphoinositide signalling, which plays a role in plant defense^19^, as well as P-type ATPases, which are crucial for ion transport, pH regulation, nutrient uptake, metal detoxification, and maintaining electrochemical gradients essential for plant growth and stress responses^20,21^. These findings are consistent with reports in other plant pan-genomes^22,23^. Core genes tended to be more highly expressed than those in the cloud and shell compartments (**Extended Data Fig. 3e**). In all polyploid oat genomes and tissues, the mean expression of genes designated to the C-genome was significantly lower than that of their A and D counterparts, with a Fisher’s Exact Test p-value of 5.46×10⁻⁴⁵ **(Supplementary Table 5, Extended Data Fig. 4a).** This confirms trends reported previously in the first genome analysis of a single oat variety^5^.

## Diversity in gene expression dynamics thin the oat pangenome

Our replicated expression data allowed us to investigate the complexity of the oat pantranscriptome. Variation occurs at multiple, intersecting levels: copy numbers of genes vary between subgenomes and across lines, and expression can differ between tissues, subgenomes and lines. To quantify transcript abundance, we mapped the RNA-seq reads of 23 oat lines to their respective genomes. Transcript levels across lines were rendered comparable by our orthologous framework. The resultant gene expression matrices are available at the Bio-Analytic Resource^24^ (https://bar.utoronto.ca/~asher/efp_oat/cgi-bin/efpWeb.cgi). Owing to the great multitude of possible patterns, we initially focused on a set of 5,965 genes that occurred in single-copy in each of the three subgenomes in 20 hexaploid *A. sativa* genomes in our panel (A:C:D 1:1:1 configuration in each), termed “60-lets”. The expression patterns of these “triads” in each genome were classified into one of seven categories based on the Euclidean distance to seven ideal expression level profiles: A-, C-, or D-dominant/suppressed, where one gene is predominantly expressed or suppressed, and a balanced category, where A, C, and D genes are equally expressed^5,25^ **(Extended Data Fig. 4b)**. Almost half (49.4%) of the 60-lets had the same classification into one of these categories across lines and were termed “stable triads”. In most cases (94% on average, with variations across tissues), the expression of stable triads was balanced among subgenomes (**Fig. 2b** and **Supplementary Table 6)**. These stable balanced triads were, for example, enriched for essential cellular functions such as vesicle trafficking, ribosome biogenesis, and protein biosynthesis and modification (**Supplementary Table 7**). In an average of 3% of the stable triads, expression was unbalanced, with the bias most often appearing in the C-genome ortholog **(Supplementary Table 6, Extended Date Fig. 4c)**.

Expression in roots, embryos and panicles tended to be more conserved across lines, while expression in leaf, internode and caryopsis tissues was more variable (**Fig 2b,c, Supplementary Table 6**). For example, HD-ZIP I/II transcription factors, which were enriched among stable genes in embryo tissue, are critical regulators in plant embryogenesis^26,27^ and stable genes in root tissue were enriched for genes involved in root formation **(Supplementary Table 8)**. This variability was further highlighted by calculating Cramer’s V matrices using the expression level categories, where we observed a more comprehensive range of values for leaf, internode, and caryopsis, indicating greater variability in these tissues compared to roots, embryo, and panicle (**Fig. 2c**). To determine how DNA sequence and expression diversity are related, we compared the matrices derived from expression data to genetic distance matrices derived from SNP data. The clustering analysis performed using these matrices showed broad agreement with genomic distance measures (**Extended Data Fig. 4d)**, suggesting that genetic factors underlie the expression patterns we observed.

When analysing stability of 60-lets not only across all lines but also across all six analysed tissues, only 13.5% of genes maintained stability. This significant reduction in overall stability indicates considerable tissue-specific differences in gene expression patterns across genotypes. This set of very conserved genes was enriched for fundamental cellular processes such as protein biosynthesis, vesicle trafficking, and RNA processing, among others **(Supplementary Table 9).** On average, 50.1% of the triads exhibited variability in their classification into one of the seven expression level categories across the 20 *A. sativa* lines, and these were termed ‘variable triads’. Nearly all these variable triads exhibited expression variability in both lines and tissues. Single-copy orthologs with expression variability were enriched for several families of transcription factors, including R2R3-MYB, ERF, NAC, and MADS/AGL (**Supplementary Table 7).**

After focusing on 60-lets, we relaxed our criteria to include genes that occurred as 1:1:1 single-copies in some genomes, but lacked the copy of one subgenome in others (1:1:0, 1:0:1, 0:1:1). We compared the expression patterns of “complete” and “incomplete” triads between 20 *A. sativa* lines to see how gene loss in the hexaploid has affected the expression of the remaining homeologs. The set of “incomplete triads” comprised 944 members. Among these, 326, 338 and 280 lacked the A, C and D gene copies, respectively. Loss of either the A or D genome copy was accompanied by a significant expression increase in the remaining D or A copy in 13.3% and 23.2% of cases, respectively. Higher expression of the C genome compensated for the loss of A or D genes in only 6% and 5.9% of cases, respectively. Similarly, if a C-derived gene was lost, compensatory up-regulation of its A or D homeologs occurred in just 4.3% and 4% of cases, respectively (**Supplementary Table 10**). The A and D genomes diverged less than 4.3 million years ago (mya) and are thus more closely related to each other than to the C genome, which split from the A/D lineage 8 mya^6^. This lower evolutionary distance suggests that A and D genes can more frequently compensate for each other than for a lost C-subgenome copy.

## Genetic diversity within the oat cellulose synthase superfamily

Oat grains contain beta-glucans, a type of polysaccharide that is known to reduce the blood LDL-cholesterol level which contributes to lower the risk of cardiovascular disease in humans^28^. One class of genes known to play a role in the biosynthesis of beta-glucans is the cellulose synthases. These enzymes are members of glucosyltransferase family 2 and build up the long chains of UDP-glucose molecules that make up both cellulose and beta-glucans. We determined and compared genic copy numbers and expression levels in the cellulose synthase and the seven cellulose synthase-like gene families in 20 *Avena sativa* lines, including a diploid (A), a tetraploid (CD), and synthetic hexaploid Amagalon. On average, we found 44.3 cellulose synthase or cellulose synthase-like genes per haploid (sub-) genome (133.2 per hexaploid genome; n=21). We observed diversity in both gene expression and copy number variation among the PanOat lines (**Fig. 2f, Extended Data Fig. 4e, Supplementary Table 11 and 12**). Moreover, we found that certain major genes contributed significantly to the gene expression of their respective subfamilies (**Supplementary Table 12, Extended Data Fig. 4f)**.

For the most important families of the cellulose synthase superfamily, CesA (cellulose synthases) and CslF (cellulose synthase-like F), which on average contributed to 44% and 26% of overall gene expression of this gene superfamily respectively, we found a moderate correlation between gene expression and genic copy number of expressed genes (tpm >=0.5; CesA: r = 0.5, p-value = 0.017; CslF: r = 0.55, p-value = 0.01). The predominant contribution by a few key genes might explain why copy number variation has a limited impact on overall expression levels.

Moreover, our analysis revealed that cultivars tend to have a higher overall expression than non-cultivars of genes encoding for cellulose synthase and cellulose synthase-like gene families **(Supplementary Table 13)**. This observation underscores the importance of following a pan-genomic approach to capture the full extent of genetic and expression diversity. The presence of major genes contributing significantly to the expression in all lines indicates that certain key regulatory elements may be conserved and critical for beta-glucan biosynthesis.

## A chromosomal inversion linked to early heading

Our chromosome-scale reference sequences made it possible to study the prevalence and impact of large structural variants in domesticated oats. The mosaic genome of hexaploid oat is the outcome of rare chromosomal rearrangements that have accumulated over evolutionary timescales^5,6^. Three chromosomal rearrangements that are polymorphic in cultivated oat and its immediate wild progenitor *A. sterilis*, were previously discovered using cytological techniques^9^. Not unexpectedly, many of these known events were identified in our genome assemblies (**Supplementary Table 14**), including (i) a translocation from chromosome 1C to 1A that has been implicated in phenological shifts associated with winter types; (ii) a translocation from 6C to 1D^29^, and (iii) a large pericentric inversion on chromosome 3C (**Fig. 1c, Extended Data Fig. 5a**). In addition, the pangenome revealed three previously unknown major structural rearrangements: an alternate reciprocal version of the 1A/1C translocation found in the *A. occidentalis* CN29555 assembly, a 2A/2C translocation, and a large 420 Mb pericentric inversion on chromosome 7D (**Fig. 3a**). The non-reciprocal 1C to 1A translocation polymorphism is present in *A. sterilis* and thus must have arisen before domestication^9^. The same applies to the 3C inversion and the pericentric inversion on chromosome 7D, whose allelic state differs between the two *A. sterilis* accessions (TN1 and TN4) in the PanOat panel (**Fig. 1c** and **Extended Data Fig. 5a, b**). Tinker et al.^30^ observed a lack of recombination in a cross that was polymorphic on most of chromosome 7D and postulated the presence of an inversion, a prediction that was borne out by our data. Although we were not able to design diagnostic markers spanning the inversion breakpoints owing to high sequence divergence in these regions, we observed two long pericentromeric haplotypes among PanOat lines that were diagnostic of the inversion. These haplotypes were identified in short-read resequencing data (5-fold coverage) of 295 oat varieties from the Collaborative Oat Research Enterprise (CORE) panel^31^, which is composed mainly of North American varieties (**Supplementary Table 15, Fig. 3b, Extended Data Fig. 6a**). Those PanOat genomes that had the ancestral state of the 7D inversion (also seen in *A. insularis*) had the less common of the two haplotypes (#1), whereas those with the derived allele, all had the more frequent haplotype (#2). A genome-wide association scan (GWAS) for heading date yielded two prominent peaks on chromosomes 7A and 7D consistent with prior genetic mapping^31^ (**Figure 3c**). Carriers of Haplotype 1 on chromosome 7D flowered on average 3.7 days earlier than those of Haplotype 2 (**Figure 3d**, **Extended Data Fig. 7b,c**). Further research is required to find out whether the inversion is associated with flowering due to suppression of recombination between multiple genes affecting early- and late-flowering haplotypes or, alternatively, if it influences heading date directly, for instance by leading to altered expression patterns of flowering time regulators or affecting the proximity of genes and regulatory factors. Three important quantitative trait locus (QTLs) are mapped to chromosome 7D involving flowering time (CO and VRN3/FT)^30^ and daylight insensitivity (Di)^32^. Further work is needed to determine the difference between the alleles in the two haplotypes.

**Figure 3:**
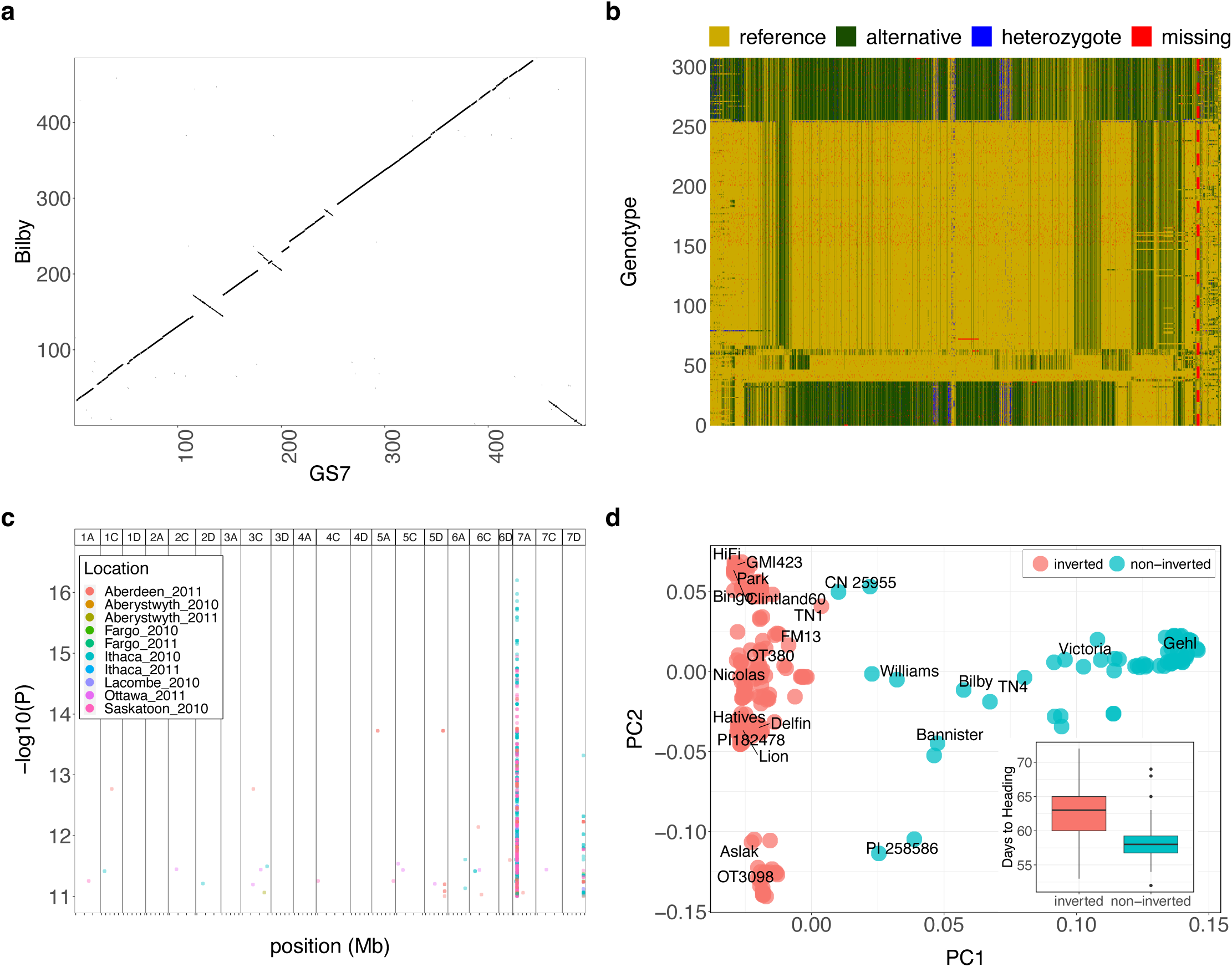
A chromosomal inversion linked to early heading. **(a)** Genome alignment of the ancestral chromosome 7D in the Australian cultivar Bilby and the inverted chromosome 7D in cultivar GS7 shows a large 450Mb pericentric inversion and a small 50 Mb distal segment that is the same in both forms. **(b)** Haplotype plot of 307 CORE and hexaploid PanOat lines sorted by predicted inversion state (top – green – ancestral and bottom – yellow – inverted), based on SNP calling against the GS7 genome. The red line marks the inversion position. **(c)** Significant kmer-GWAS results for ten locations in 2010 and 2011. Two significant peaks are visible on chromosomes 7A and 7D. **(d)** PCA plot of CORE31 lines using SNPs at the distal end of chromosome 7D (400-495Mb) overlaid with PanOat assemblies (**Supplementary Table 1**). The boxplot shows the difference in heading time in the Ithaca 2010 field trial between inverted and non–inverted lines. PI 258586 = A. byzantina PI258586, HdA = Hâtives des Alpes, Nicolas = AAC Nicolas, Morgan = AC Morgan, TN1 = A. sterilis TN1, TN4 = A. sterilis TN4, and CN 25955 = A. occidentalis CN 25955.

## The hidden legacy of mutation breeding

One of the previously unknown translocations discovered in this study was a homeologous exchange of the short arms of chromosomes 2A and 2C in the Australian variety Bannister (**Fig. 1C Fig. 4a**). This is unlikely to be an assembly artefact because an intense interchromosomal signal was observed in Hi-C data of Bannister mapped to the genome of GMI423, an accession with the standard karyotype (**Extended Data Fig. 7b**). Sequence data from three recombinant inbred lines (RILs) derived from a cross between Bannister and non-translocated Williams revealed whole chromosome arm deletions and compensating duplications on chromosomes 2A and 2C, which are most readily explained by anomalous meiosis in heterozygotes (**Extended Data Fig. 7d**). Furthermore, we designed diagnostic PCR marker assays spanning the translocation breakpoints to confirm the presence of the 2A and 2C translocations **(Extended Data Fig. 7c**). Finally, C-banding in Bannister confirmed the presence of recombined 2A and 2C chromosomes with pronounced differences in heterochromatin content between long and short arms (**Fig. 4a**).

**Figure 4:**
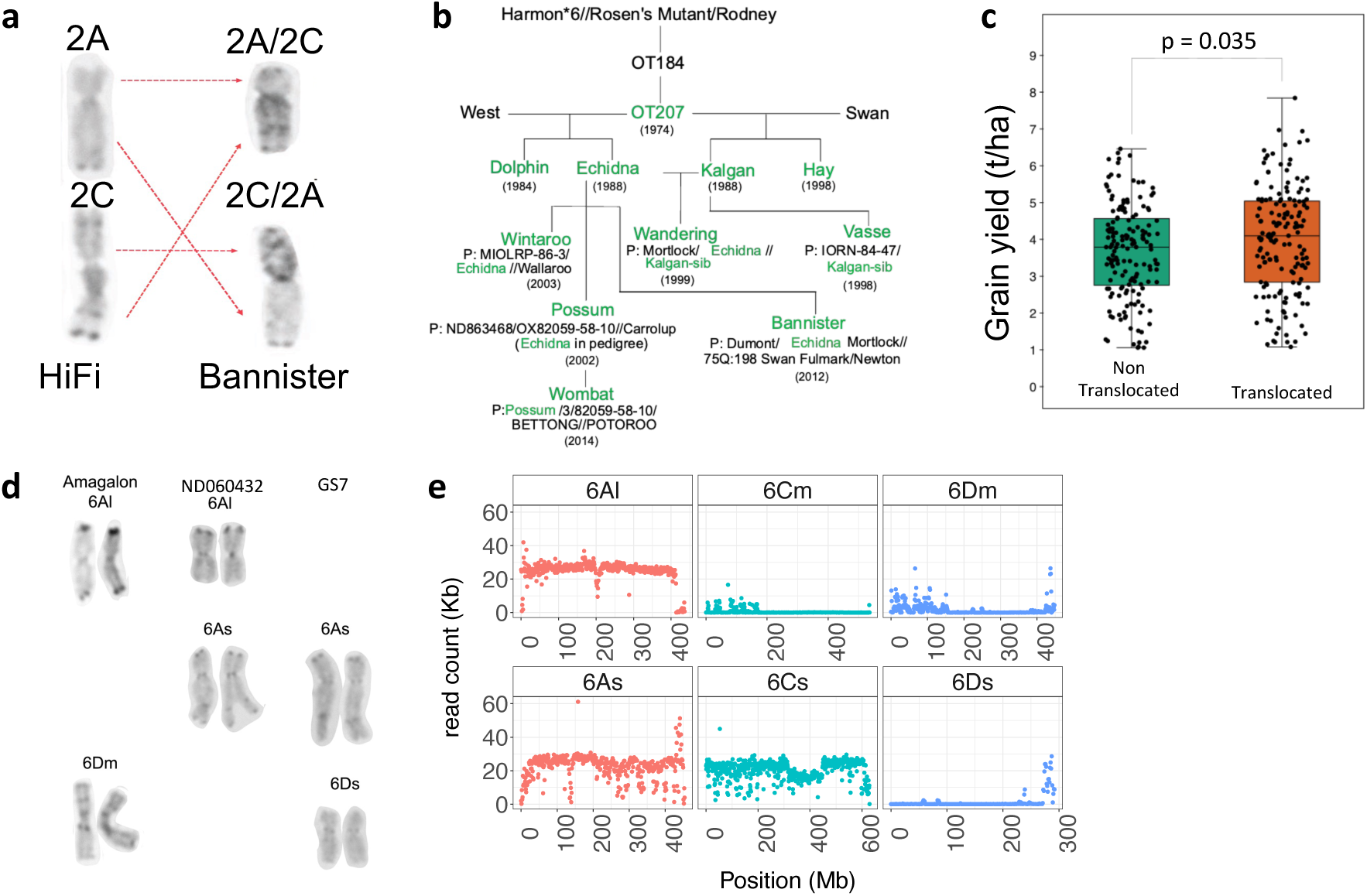
Chromosomal translocations and homeologous sequence exchanges. **(a)** Karyotype analysis comparing chromosomes 2A and 2C between a non-translocated variety (HiFi) and a translocated one (Bannister). **(b)** Origin of the 2A/2C translocation and inheritance in Canadian and Australian oat varieties. **(c)** Yield advantage of varieties carrying the 2A/2C translocation in Australia’s National Variety Trials. **(d)** Karyotypes of Amagalon, A. sativa - ND060432 and A. sativa – GS7. The karyotypes show the lack of chromosome 6D in ND060432 and its substitution with chromosome 6A from Amagalon. **(e)** Read mapping counts of whole-genome resequencing data from ND060432 showing the substitution of A. sativa 6D with Amagalon 6A. Reads were mapped to a hybrid reference combining Amagalon (top) and A. sativa GS7 (bottom).

To measure the frequency of this reciprocal translocation in a broader set of oat germplasm, we used the translocation diagnostic markers to genotype 564 accessions from 41 countries around the world and found 17 (3 %) that carried the translocation (**Supplementary Table 16**). Carriers were mainly of Australian origin and included cultivars released since the last quarter of the 20^th^ century. Among them were the ‘Dolphin’, ‘Echidna’, ‘Kalgan’, and ‘Hay’ as well as current highly successful varieties ‘Bannister’, ‘Possum’ and ‘Wombat’. These varieties share a common parent in OT207, a mutation breeding line created in 1970 using fast neutron-irradiation^33^. OT207 carries the 2A/2C translocation and the *Dw6* mutant allele on chromosome 6D^34^, conferring the semi-dwarf growth habit, but none of its parental varieties do (**Fig. 4b**). As expected, *Dw6* is not linked to the 2A/2C translocation and occurs also in the latter’s absence, e.g. in ‘Bilby’. It is thus most likely that the translocation is an unintended side effect of the mutagenesis. However, it is intriguing that the 2A/2C translocation has become so frequent in Australian varieties. We speculated that it may have adaptive advantages in the Australian environments. To test this hypothesis, we obtained grain yield data from the Australian National Variety Test for the years between 2017 to 2022, which represented 19 to 31 trials per year across all of Australia (158 field trials). Varieties carrying this translocation are among the highest-yielding varieties of Australia and, as a group, significantly (t-test, p = 0.03) outyielded their non-translocated counterparts in 131 field trials^31^ (**Fig. 4c**, mean yields of 3,991kg/ha vs 3,667kg/ha).

To study the effect of the translocation, we analysed a population of recombinant inbred lines (RILs) derived from a cross between Bannister, which carries the translocation, and non-translocated Williams, which does not. Among 285 RILs, we identified 10 with chromosomal arm deletion, these showed harmful agronomic traits such as reduced fertility and grain shattering. Sequencing the genomes of three of these RILs from the above 10 lines confirmed whole chromosome arm deletions and compensating duplications on chromosomes 2A and 2C, which are most readily explained by aberrant meiosis as heterozygotes.

Using the Bannister x Williams population, previous mapping results^34^ and the genome sequences of both parents to fine-map this locus, we narrowed down the QTL for plant height to a 554 kb interval on chromosome 6D. Three members of the PanOat panel, Bilby, Bannister and OT3098, are semidwarfs and share a common haplotype in the *Dw6* interval. This haplotype comprises a 41 kb large insertion containing nine genes, one of which encodes a fatty acid hydroxylase (**Supplementary Table 17**). Further functional studies are required to test if this gene is casual to the *Dw6* phenotype.

## Chromosomal rearrangements in oat breeding germplasm

Our pangenome panel includes ‘Amagalon’, a synthetic hexaploid oat that was derived from a cross between diploid *A. longiglumis* (A_l_A_l_; CW-57) and tetraploid *A. magna* (C_m_C_m_D_m_D_m_; CI8330), a close relative of *A. insularis* (C_i_C_i_D_i_D_i_) – the putative donor of the CD genome in hexaploid oat^35^. A source of disease resistance genes^36^, Amagalon and a derived cultivar HiFi have been widely used in US and Canadian oat breeding. The Amagalon assembly revealed several chromosomal translocations relative to other oat varieties (**Extended Data Fig. 5c)**. To determine if these structural variants may have affected inheritance in Amagalon-derived breeding lines, we analysed WGS and GBS data and identified numerous regions in which read depth in 1 Mb windows deviated from the genome-wide average (**Extended Data Fig. 8** and **Extended Data Fig. 9**). The sizes of these regions varied from several Mb to whole chromosomes, where the read depth halved, doubled, or declined to zero. We found four lines from the North Dakota State University (ND) breeding program in our WGS panel (**Supplementary Table 15**) that showed a drop in read depth on chromosome 6D and a concomitant increase on chromosome 6A. The pedigree of these lines suggested that some of them had Amagalon in their pedigree (https://oat.triticeaetoolbox.org/). We analysed the karyotypes of Amagalon, ND060432 (chromosome 6D_s_ missing) and GS7 (chromosome 6D_s_ present) and confirmed the presence of two 6A chromosomes in ND060432, 6A_s_ and 6A_l_ (**Fig. 4d**). When the sequence data of the four lines were aligned to a reference combining Amagalon and the variety GS7, few reads mapped to either of the 6D_s_ reference chromosomes, however, reads abundantly mapped to both 6A_s_ and 6A_l_ chromosomes coming from GS7 and Amagalon, respectively (**Fig 4e**). This suggests that the original cross involving Amagalon resulted in the *A. sativa* 6D_s_ chromosome being replaced with a 6A_l_ chromosome, which is ultimately derived from *A. longiglumis.* Owing to the low resolution of C-banding, it is unlikely that this event could have been discovered without genome sequencing and pan-genome analysis. Since the ND lines were included in the CORE panel for their promising agronomic performance, exchange between homeologous chromosomes seems to be tolerated and may be exploited in future breeding programs.

To gauge the extent to which SVs impact oat breeding, We genotyped and analysed recombination and segregation patterns in 13 bi-parental populations to assess the impact of SVs in an active oat breeding program. Numerous crosses exhibited segregation distortion as well as non-linear or suppressed recombination within or between chromosomes (**Extended Data Fig. 10a**). These include regions on chromosomes 1A, 1C, 3C, 4C, and 7D, where inversions and translocations identified in this study and the companion global oat genomic diversity analysis (**Bekele *et al*., 2024**) were confirmed. We analysed and karyotyped two half-sib crosses in greater detail (**Extended Data Fig. 10 b to f**). One cross exhibited pseudo linkage between chromosomes 1A and 1C associated with a 1A/1C reciprocal translocation heterozygote, as well as recombination suppression on chromosome 7D associated with inversion heterozygotes. The other showed recombination suppression on chromosome 3C, also associated with known chromosome inversion. Signatures of other SV heterozygotes were observed, including a potential translocation between chromosomes 1D and 7C. The companion study (**Bekele, *et al*., 2024**) introduces a SNP-based in-silico karyotyping method that could help breeders avoid crosses with segregation irregularities, to design genomic selection strategies to reduce linkage drag, or to leverage suppressed recombination to preserve blocks of adaptive alleles.

## Discussion

The oat pangenome and its detailed analysis reported here should accelerate the adoption of genomic methods in oat research and breeding. Multiple reference genome sequences, an extensive gene expression atlas, and resequencing data of a well-characterised diversity panel greatly expand the genomic resources available to oat researchers and will have an immediate impact on genomics-assisted breeding. As in other crops, the full extent of structural variation in elite varieties has been unknown in oats. In tracing the origin and impact of a recent homeologous recombination event, the 2A/2C translocation, we encountered a situation that is analogous to one described by Jayakodi et al. in barley^37^. These authors discovered a 141-Mb paracentric inversion, which was most likely induced by irradiation and spread due to a founder effect because the mutant variety had a desired semi-dwarf growth habit. This supports the notion that mutation breeding in the 20^th^ century resulted in positive changes to agronomically relevant traits, but also led to cryptic chromosomal aberrations that continue to influence crop improvement in surprising ways. Cytological evidence^38^ leads us to expect that more such events will be discovered in oat germplasm.

A pangenome is an indispensable resource for applications in marker discovery, genetic mapping, and molecular breeding. A map of large structural variants will help breeders interpret segregation patterns in crosses and guide the selection of parents. We also hope that the resources we have built will stimulate oat research beyond translational applications and will motivate more scientists to make oat a major research focus. There are still many open research questions in wild and domesticated oat that remain unanswered, including the genetic basis of the domestication syndrome in oat. Its key components are the loss of grain shattering, reduction of the geniculate awn, loss of lemma pubescence, and husk adherence^39^, as well as loss of dormancy and changes in photoperiod or vernalization response. Some encouraging progress in the latter trait has been made thanks to the genome sequence of a hulless oat^6^. The next steps in oat genomics should be to expand the pangenome of cultivated oats and work towards a genus-wide pangenome of *Avena* comprising all the approximately ∼30 species in the genus. The latter effort would underpin inquiries into structural genome evolution, which have so far been limited to the hexaploid and its immediate progenitors.

## Acknowledgements

We thank Prof. Jörg-Peter Schnitzler from Helmholtz Munich for the pictures of oat plants. We would like to acknowledge Sirpa Moisander, Santeri Kankanpää, for their assistance in RNA and DNA extractions; Auli Kedonperä, Marjo Segerstedt, Marja Jalli and Anne Nissinen from the Natural Resources Institute Finland (LUKE) for assistance in growing oats under greenhouse conditions and for photo documentation; Ed Wilcox from the DNA sequencing center at Brigham Young University for support and expertise with PacBio sequencing . M.M., M.S. and M.H.H. were supported by a grant from the German Ministry of Food and Agriculture (BMEL, FUGE, FKZ: 28AIN02C20). W.A.B. and N.A.T. acknowledge co-funding by Genome Quebec through a project entitled “Targeted and useful genomics for Barley and Oat (Tugboat). N.K. was supported by the European Research Council (ERC) under the European Union’s Horizon 2020 Research and Innovation programme (Grant Agreement No 101116452, ERC Starting Grant project ‘RESIST’). E.P. and F.J.C. were supported by grant [PID2022-142574OB-I00] funded by MICIU/AEI /10.13039/501100011033, FEDER, UE, and Junta de Andalucia [QUAL21_023 IAS]. S. Marmon and N. Sirijovski acknowledge funding from the Swedish Foundation for Strategic Research (ScanOats: IRC15-0068) and the two industrial partners of the ScanOats Research Center (Lantmännen and Oatly AB); support from the Swedish National Genomics Infrastructure funded by the Science for Life Laboratory, the Knut and Alice Wallenberg Foundation and the Swedish Research Council; the SNIC/Uppsala Multidisciplinary Center for Advanced Computational Science for providing assistance with massively parallel sequencing and access to the UPPMAX computational infrastructure; and Johan Bentzer at Lund University for data management support. C.J.H. and T.L. were supported by BBSRC grants BBS/E/IB/230001, BBS/E/W/0012843 and BB/S008195/1. JIS was supported by Beatriz Galindo Program (BEAGAL18/00115) grant, PID2021-123718OB-I00, funded by MCIN/AEI/10.13039/501100011033 and by ‘‘ERDF A way of making Europe,’’ CEX2020-000999-S and Severo Ochoa Program for Centres of Excellence in R&D from the Agencia Estatal de Investigación of Spain, Grant SEV-2016-0672. K.T.N. acknowledges the technical support provided by Sudhakar Pandurangan, and the funding provided by the Prairie Oat Breeding Consortium. We thank Lars Paulin and the DNA Sequencing and Genomics Laboratory, Institute of Biotechnology, University of Helsinki, for whole genome sequencing of ‘Aslak’; and Natural Resources Institute Finland (LUKE) for financial support. We acknowledge the Finnish Functional Genomics Centre core facility, Turku Bioscience for RNA sequencing. The Functional Genomics Center Zurich (Switzerland) is acknowledged for their contribution to whole genome sequencing of Hâtives des Alpes accession. Australia GRDC (UMU2003-002RTX) and Western Australia Oat Industry Partnership and Department of Primary Industry and Regional Development (POIGP/01) provided funds for C.L. Dr Allan Rattey from InterGrain provided the Bannister/Williams RIL population; the Australia National Variety Test Program provided long term yield data of Australian oat varieties. T.Langdon was supported by BBSRC award BB/S008195/1 ‘Oat domestication’. Funding for the sequencing of *A. sativa* cv. GS7 came from NSF award ABR-PG 1444575. This research was supported by USDA Agricultural Research Service project numbers 3060-21000-046-000D, 2030-21000-056-000D, 2050-21000-038-000D, 8062-21000-053-000D. Mention of trade names or commercial products in this publication is solely for the purpose of providing specific information and does not imply recommendation or endorsement by the U.S. Department of Agriculture. USDA is an equal opportunity provider and employer.

## Author contributions

Selection of genotypes: W.A.B., L.B., B.B., F.J.C., S.R.E., K.E.K., Y.-B.F., M.H.H., C.J.H., J.I.S., J.-L.J., E.N.J., T.Langdon, C.L., M.M., P.J.M, K.T.N., E.P.-G., E.P., A.R., J.S., N.Sirijovski, B.S., N.A.T., S.Viitala, S.W., W.Y. Genome sequencing: P.A., W.A.B., L.B., O.B., H.S.C., Y.C., D.C., V.D., S.R.E., K.E.K., Y.-B.F., R.G., A.H., C.J.H., A.I., J.-L.J., R.K., M.K., T.Langdon, C.L., S.Marmon, P.J.M, K.T.N., E.P.-G., A.R., R.W.R., J.A.S., A.H.S., J.S., M.-Singh, N.Sirijovski, N.Stein, B.S., N.A.T., S.W., P.W., A.J.W., X.-Q.Z., Z.Z. Sequence assembly: R.A., W.A.B., H.S.C., Y.C., H.H., M.M., P.J.M, K.T.N., B.S. Transcriptome sequencing and analysis: R.A., L.B., O.B., A.Fenn, J.D.F., R.G., N.K., T.Langdon, T.Lux, V.M., M.M., P.J.M, R.S.N., A.H.S. Annotation: H.G., G.H., N.K., T.Lux, V.M., K.F.X.M., M.Spannagl Analysis and interpretation of structural variants: T.T.A., R.A., W.A.B., B.C., V.D., T.H., E.N.J., Y.J., C.L., R.S.N., N.A.T., P.W., C.P.W., X.-Q.Z., G.Z. Data management and submission: R.A., L.B., A.Fiebig, M.M., S.Michel, R.S.N., A.P., N.J.P., T.Z.S., J.S., M.-Singh, S.Viitala, S.Vronces, E.Y., Z.Z. Writing: R.A., W.A.B., L.B., O.B., V.D., Y.-B.F., J.-L.J., N.K., C.L., T.Lux, M.M., A.H.S., T.Z.S., M.Spannagl, N.A.T., X.-Q.Z., G.Z. Coordination: W.A.B., L.B., O.B., J.D.F., R.G., N.K., C.L., M.M., P.J.M, A.H.S., T.Z.S., J.S., M.Spannagl, N.A.T. All authors read and commented on the manuscript.

## List of Supplementary items

**Supplementary Table 1:** Assembly statistics and genotype characteristics

**Supplementary Table 2:** Gene expression statistics of the 23 oat lines with RNAseq-data

**Supplementary Table 3:** Summary statistics of the gene annotation and BUSCO assessment

**Supplementary Table 4:** Functional enrichment analysis results of the core, cloud and shell genes

**Supplementary Table 5:** Mann-Whitney U test results for subgenome comparisons across tissues and lines and summary statistics of significant subgenome expression comparisons

**Supplementary Table 6:** Amount of stable and variable triads in the 20 *A. sativa* lines in percent

**Supplementary Table 7:** Functional enrichment analysis results of single-copy genes across expression level categories

**Supplementary Table 8:** Functional enrichment analysis of single-copy genes with stable expression in root, panicle and embryo and variable expression in leaf, caryopsis and internodes

**Supplementary Table 9:** Functional enrichment analysis results of single-copy genes with consistently stable expression across oat lines and tissues

**Supplementary Table 10:** Expression statistics from the comparison of complete triads and triads missing one member.

**Supplementary Table 11:** Cellulose synthase and cellulose synthase-like gene families in the studied lines of the oat pangenome

**Supplementary Table 12**: Gene expression statistics of the cellulose synthase and cellulose synthase-gene families in caryopsis tissue in PanOat

**Supplementary Table 13:** Comparison of gene expression between cultivars and non-cultivars across cellulose synthase and cellulose synthase-like gene families, based on z-scores of TPM values.

**Supplementary Table 14:** Large structural variation status in PanOat assemblies

**Supplementary Table 15:** 295 CORE lines sequenced at 5x and used for kmerGWAS and SNP calling

**Supplementary Table 16:** World-wide oat germplasm collection in Western Crop Genetics Alliance at Murdoch University

**Supplementary Table 17:** Assorted small tables for 2A/2C part

**Supplementary Table 18:** List of accessions used for global oat diversity PCA (Figure 1a and Extended Data Fig. 5a)

**Supplementary Table 19:** PanOat ENA project IDs for Pacbio, Illumina, HiC, RNA-Seq and Iso-Seq raw data and complete pseudomolecule assemblies

**Supplementary Table 20:** RNAseq read numbers and Bio Sample ID for each tissue and replicate.

**Supplementary Table S21:** Functional annotation of core single-copy genes performed with Mercator.

**Supplementary Table S22:** Details on 13 populations from a working oat breeding program at AAFC, Canada.

## Extended Data Figure legends

**Extended Data Fig. 1:**
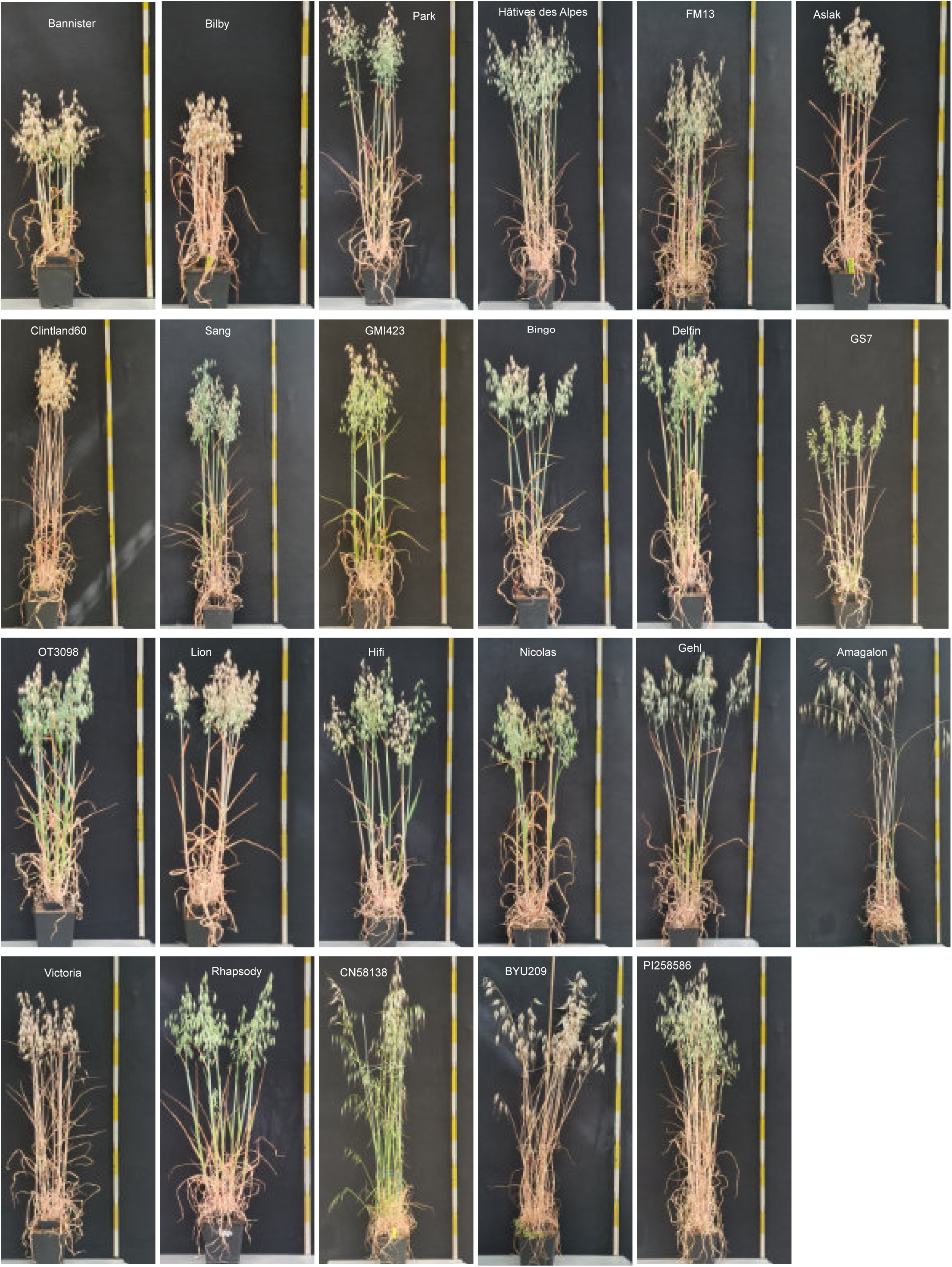
Images of the representative individual of the 23 oat genotypes for which RNA-seq data was generated.

**Extended Data Fig. 2:**
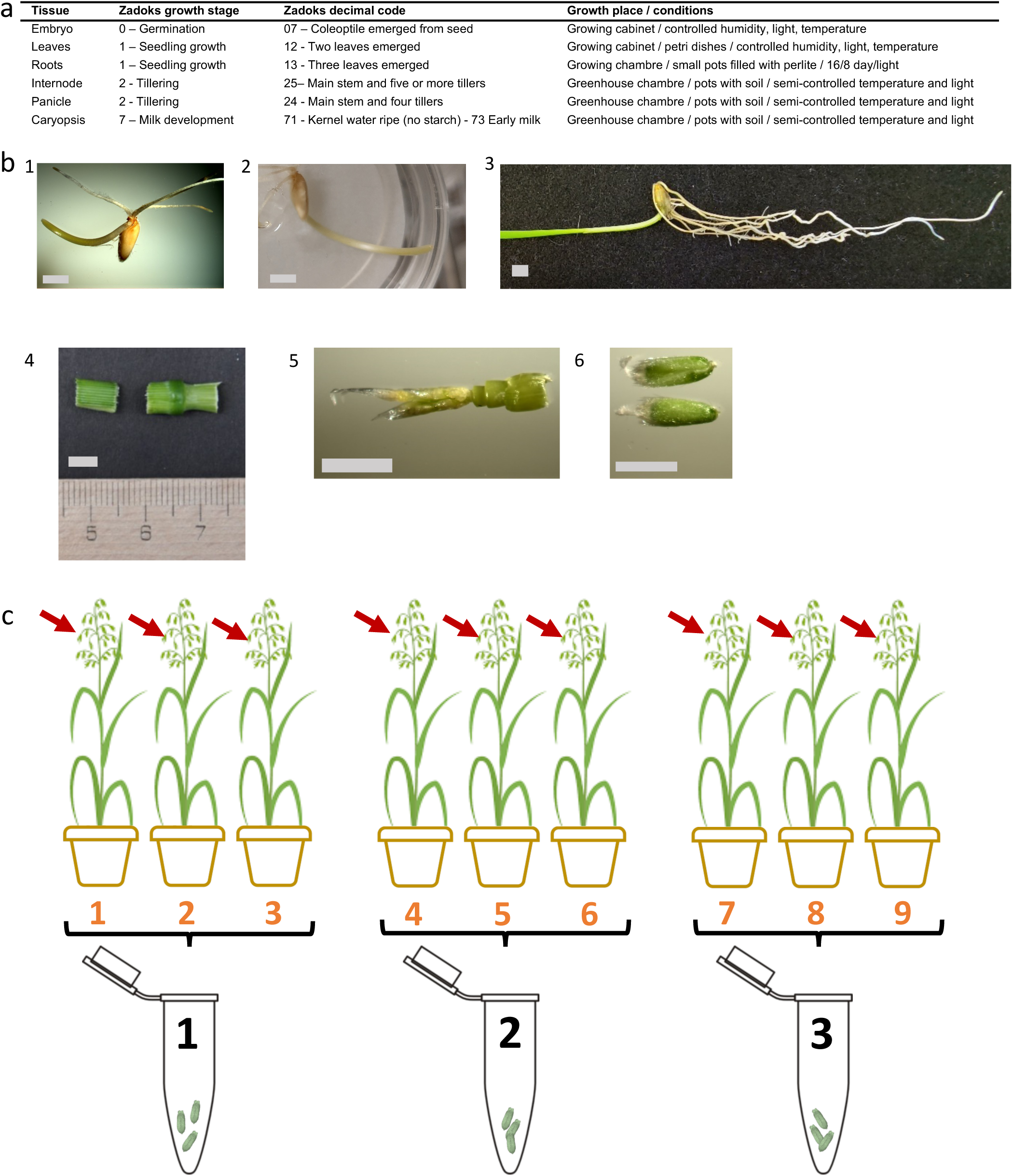
Tissue sampling and RNA extraction for transcriptome sequencing. **(a)** List of oat tissues dissected from 24 members of the PanOat panel according to defined growth stages in cereals (Zadoks scale). **(b)** Representative photos of sampled tissues in selected PanOat genotypes: 1. Germinating seed (Delfin), 2. seedling leaf (Aslak), 3. seedling root (OT3098) 4. stem (Bilby), 5. developing panicle (GMI423) and 6. immature seed (Aslak). Gray bars equel 0.5 cm. **(c)** Sampling scheme used to collect all tissues for RNAseq and IsoSeq sequencing.

**Extended Data Fig. 3:**
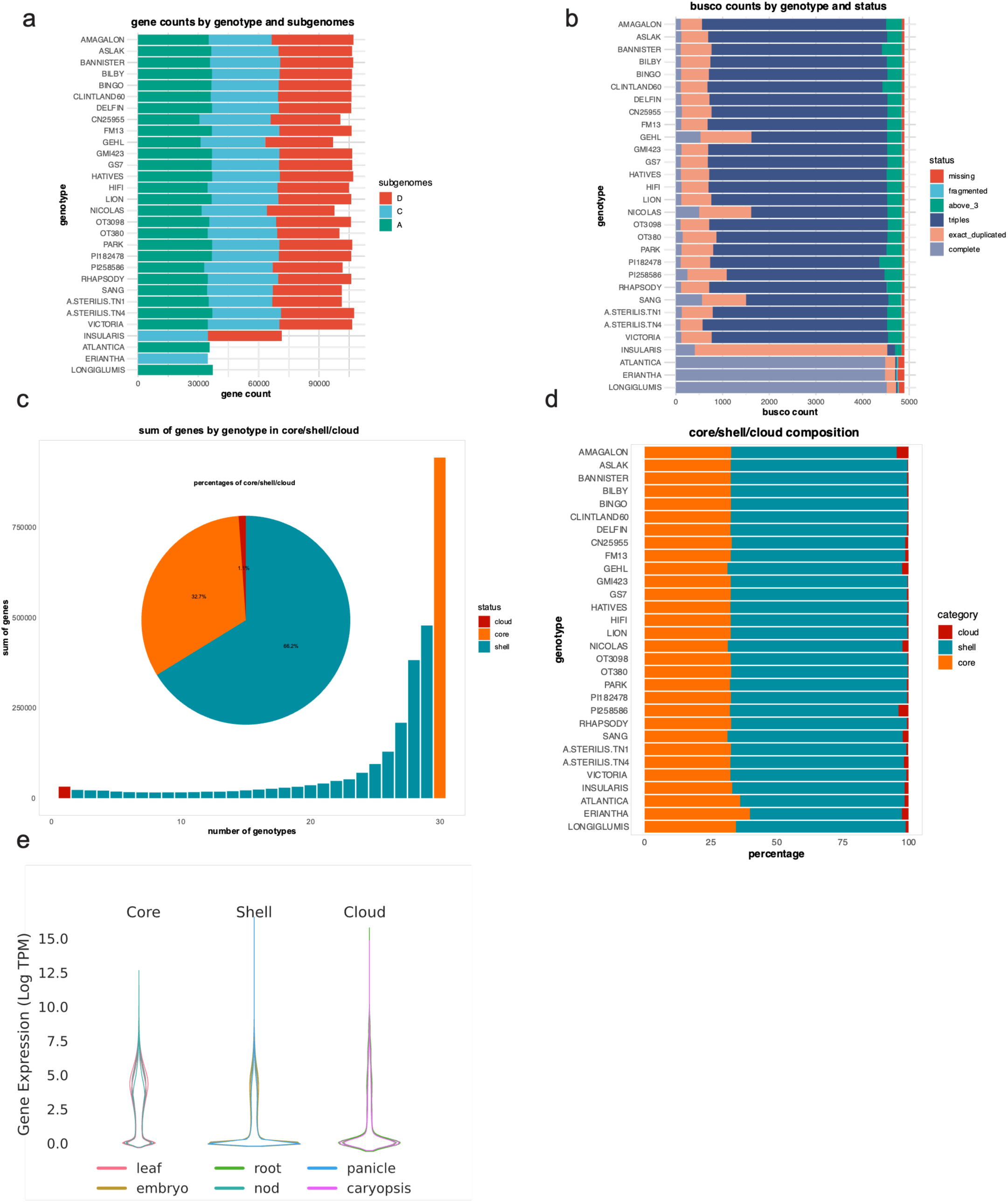
Compartments of oat pangenome. **(a)** number of genes predicted for the respective oat line, detailed by subgenome. **(b)** completeness of gene predictions assessed by BUSCO counts. **(c)** the bar chart shows the number of genes (y-axis) found in orthologous groups consisting of one oat line (cloud; red), twotwentynine oat lines (shell; turquoise) or thirty oat lines (core; orange) (x-axis). The pie chart provides the overall ratios of genes in the core-, shell- and cloud-genome categories. **(d)** ratios of genes in the core-, shell- and cloud-genome categories detailed for each oat line. **(e)** Gene expression counts as Log TPM values for the core-, shell- and cloud-genes and six different tissues.

**Extended Data Fig. 4:**
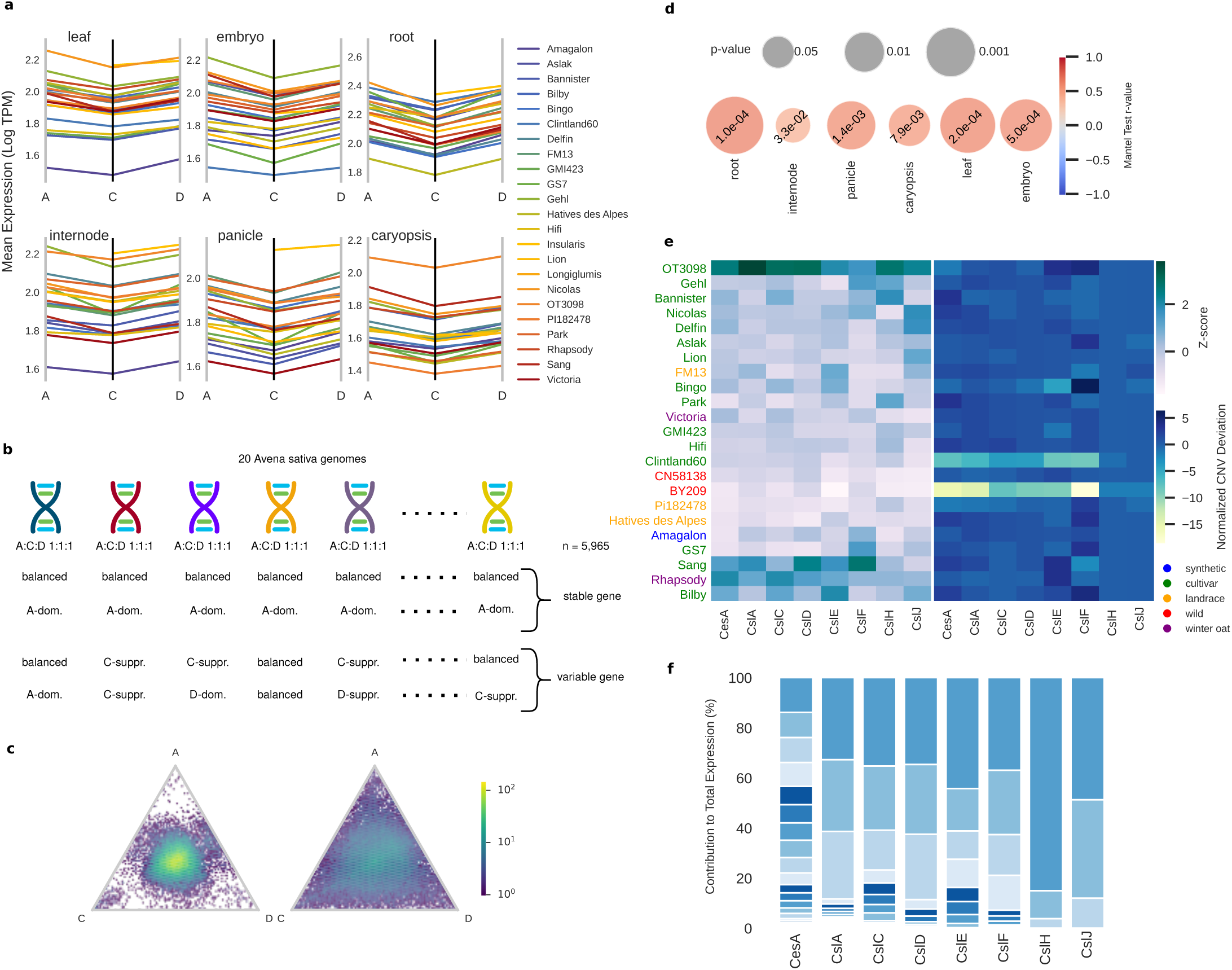
Expression diversity in the oat pangenome. **(a)** Parallel coordinates plot showing the mean expression of each of the A, C, and D subgenome in six tissues of 22 oat genomes, including 20 A. sativa genomes, synthetic Amagalon, and tetraploid BYU209 (A. insularis) in six tissues. **(b)** Mantel test p-values comparing genetic diversity based on expression level categories across six tissues and genetic distance based on SNP data in 20 A. sativa lines. **(c)** Expression level categories were identified in 20 A. sativa lines utilizing 5,294 common single-copy orthologs. Triads were classified as stable or variable based on whether all pangenome lines shared the same expression level category. **(d)** Sstable and variable genes in panicle tissue. 52.8% of triads were stable of which 50.4% were balanced and 47.2% were variable. **(e)** Gene expression and copy number variation of the cellulose synthase and cellulose synthase-like gene families in the 23 PanOat lines, that were RNA-sequenced **(f)** Stacked bar-plot of the relative contribution of individual genes to the total expression of each of the cellulose synthase and cellulose synthase-like gene families exemplified using the line “OT3098”.

**Extended Data Fig. 5:**
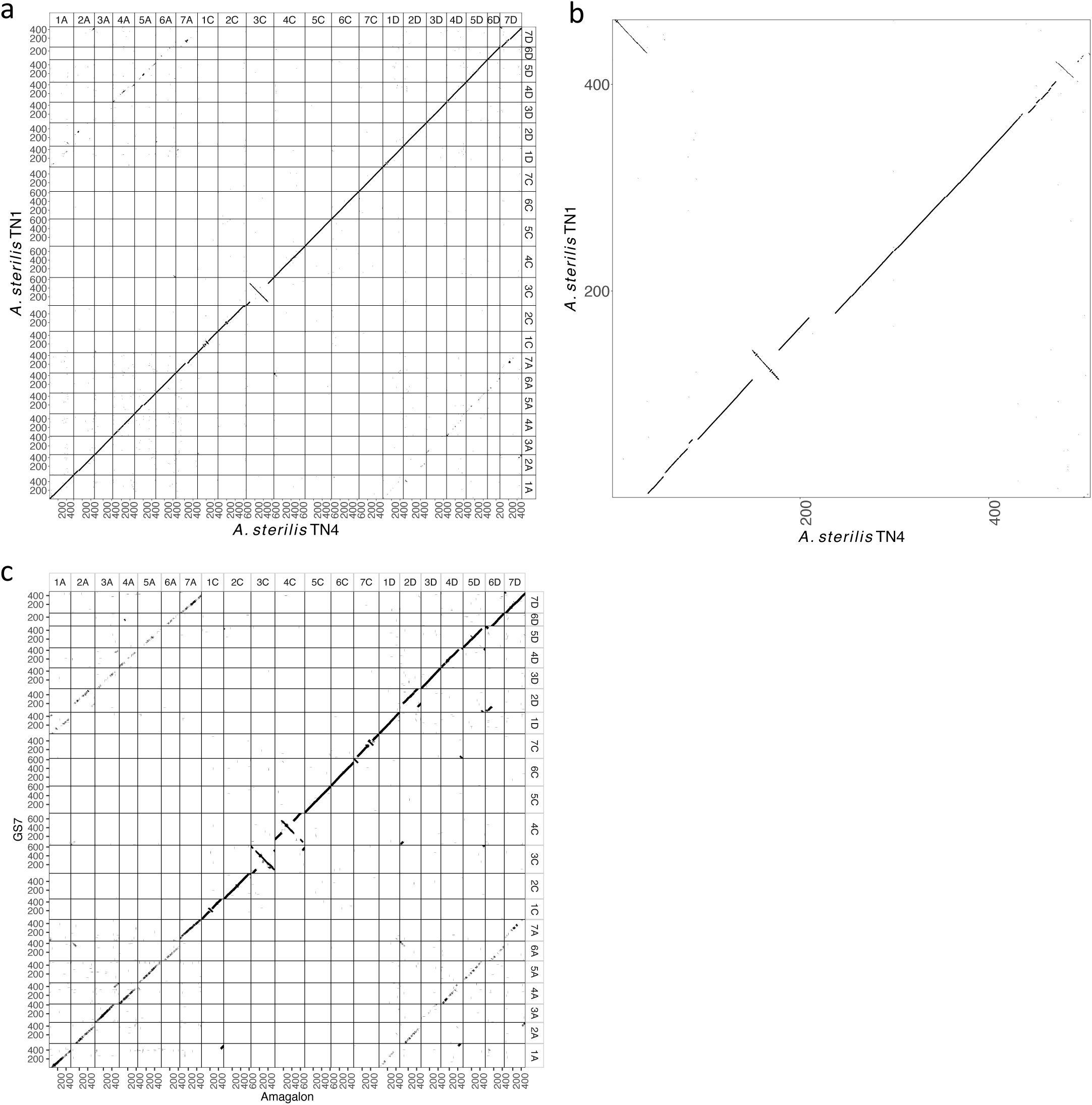
Genome alignment between A. sterilis TN1 and A. sterilis TN4 and between GS7 and Amagalon. **(a)** Alignment of all chromsomes showing two large inversions on chromosomes 3C and 7D. **(b)** Alignment of chromosome 7D showing the inversion in more detail. **(c)** Alignment of all chromosomes between GS7 and Amagalon.

**Extended Data Fig. 6.**
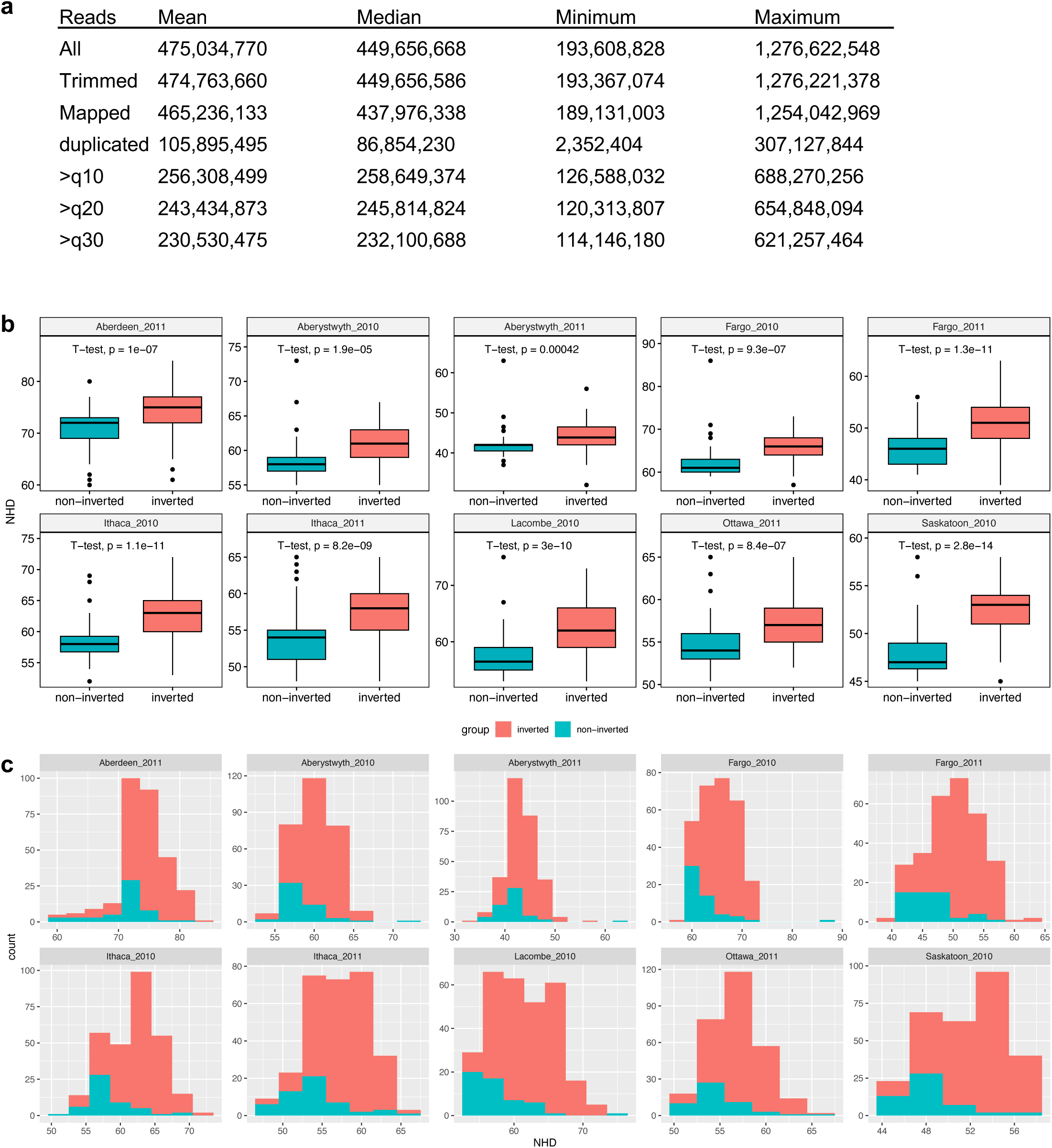
Distribution of heading date phenotype in the CORE panel in 10 environments. **(a)** WGS sequencing and mapping statistics for 295 CORE genotypes. **(b)** Boxplots and **(c)** histograms of heading date measurements (NHD) grouped by the allelic state of the inversion on chromosome 7D as inferred by PCA analysis. A t-test was used to calculate p-values.

**Extended Data Fig. 7:**
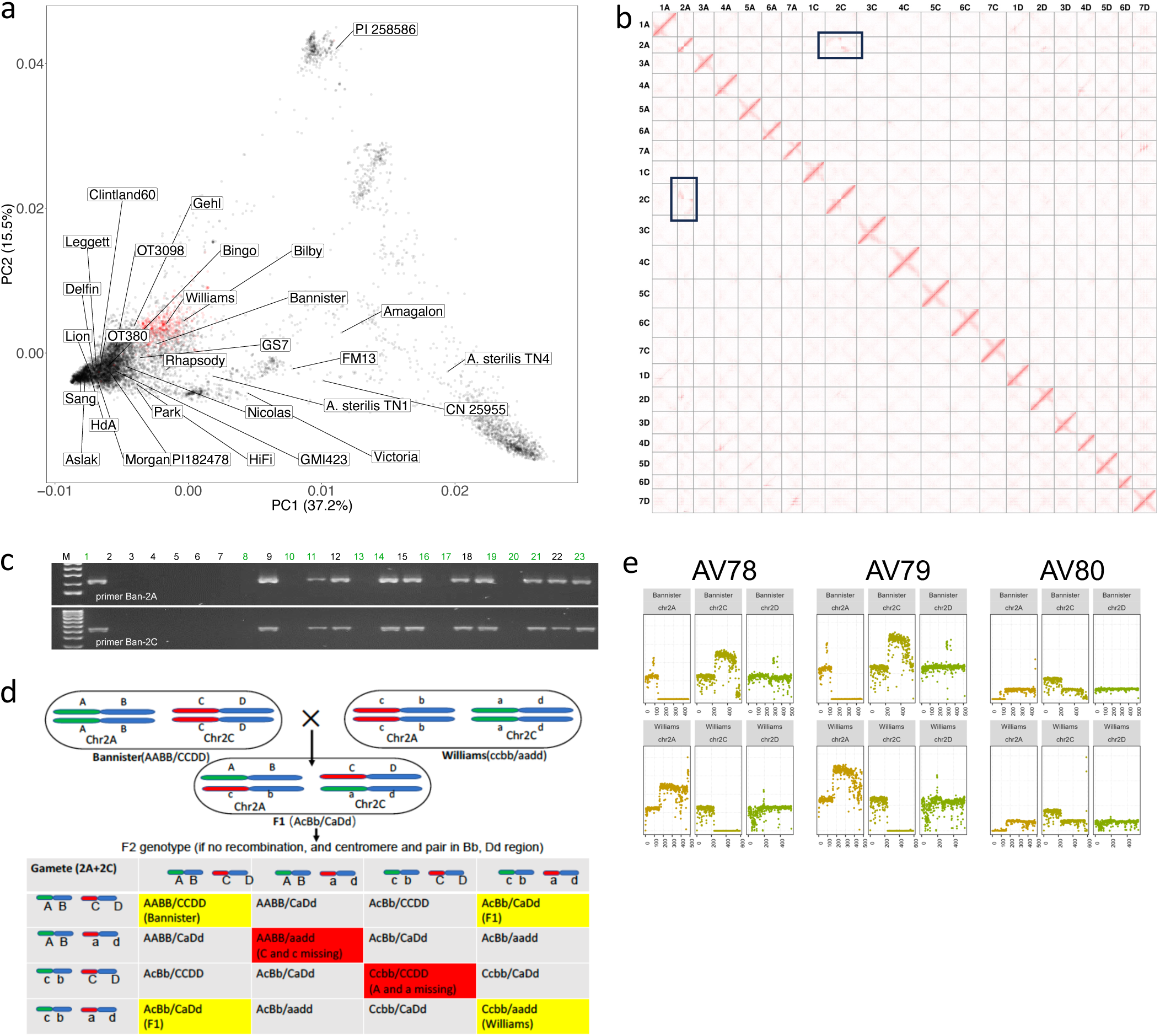
Legacy of mutation breeding in Australian oat. **(a)** PCA plot of 9,174 hexaploid oats including A .sativa, A. byzantina and A. sterilis overlaid with the positions of the 29 hexaploid PanOat lines. Australian accessions are highlighted in red. PI 258586 = A. byzantina PI 258586, HdA = Hatives des Alpes and CN 25955 = A. fatua CN 25955. **(b)** Contact matrix of the Bannister Hi-C data when aligned to the non-translocated reference GMI423. Off-diagonal signals on chromosomes 2A and 2C are due to the reciprocal translocation between these chromosomes in Bannister. **(c)** Diagnostic PCR assay to detect both translocation break points on chromosomes 2A and 2C. **(d)** Types of meiosis and synapsis in F_1_ plants and their F_2_ progenies. **(e)** Deletion and compensatory duplication of whole chromosome arms in three Bannister x Williams RILs. WGS reads were mapped to the GS7 reference genome and reads were counted in 1 Mb bins. *In panels a PI 258586 = A. byzantina PI258586, HdA = Hâtives des Alpes, Nicolas = AAC Nicolas, Morgan = AC Morgan, and CN 25955 = A. occidentalis CN 25955.

**Extended Data Fig. 8:**
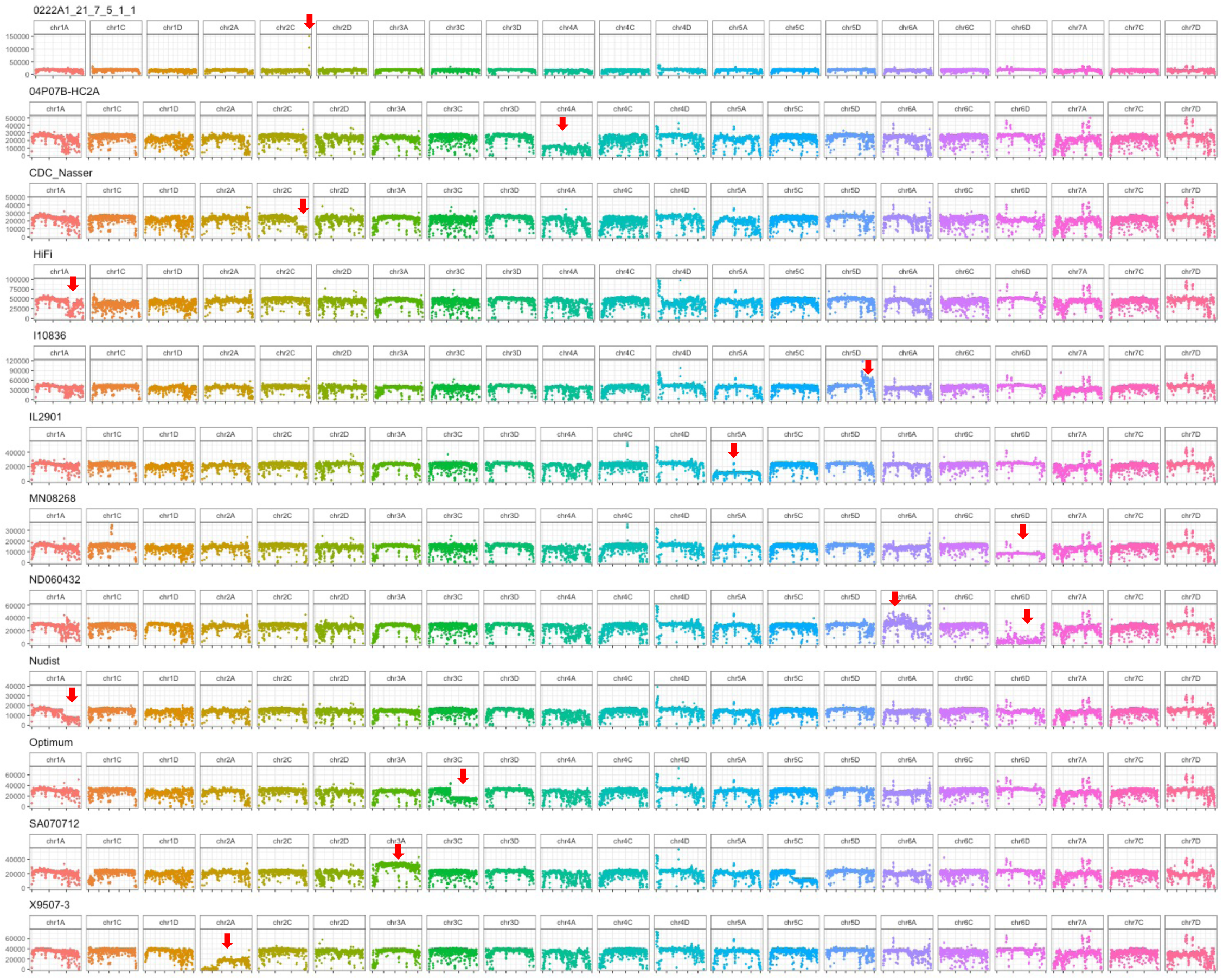
Chromosomal anomalies in representative CORE lines. WGS reads were aligned to the GS7 reference genome and read counts were aggregated in 1 Mb windows. Each row shows one genotype (**Supplementary Table 15)**. Red arrows mark SVs. At least one chromosome in each genotype is affected by large SVs, which are most likely deletions, duplications or homeologous exchanges. A detailed example is described in Fig. 3.

**Extended Data Fig. 9:**
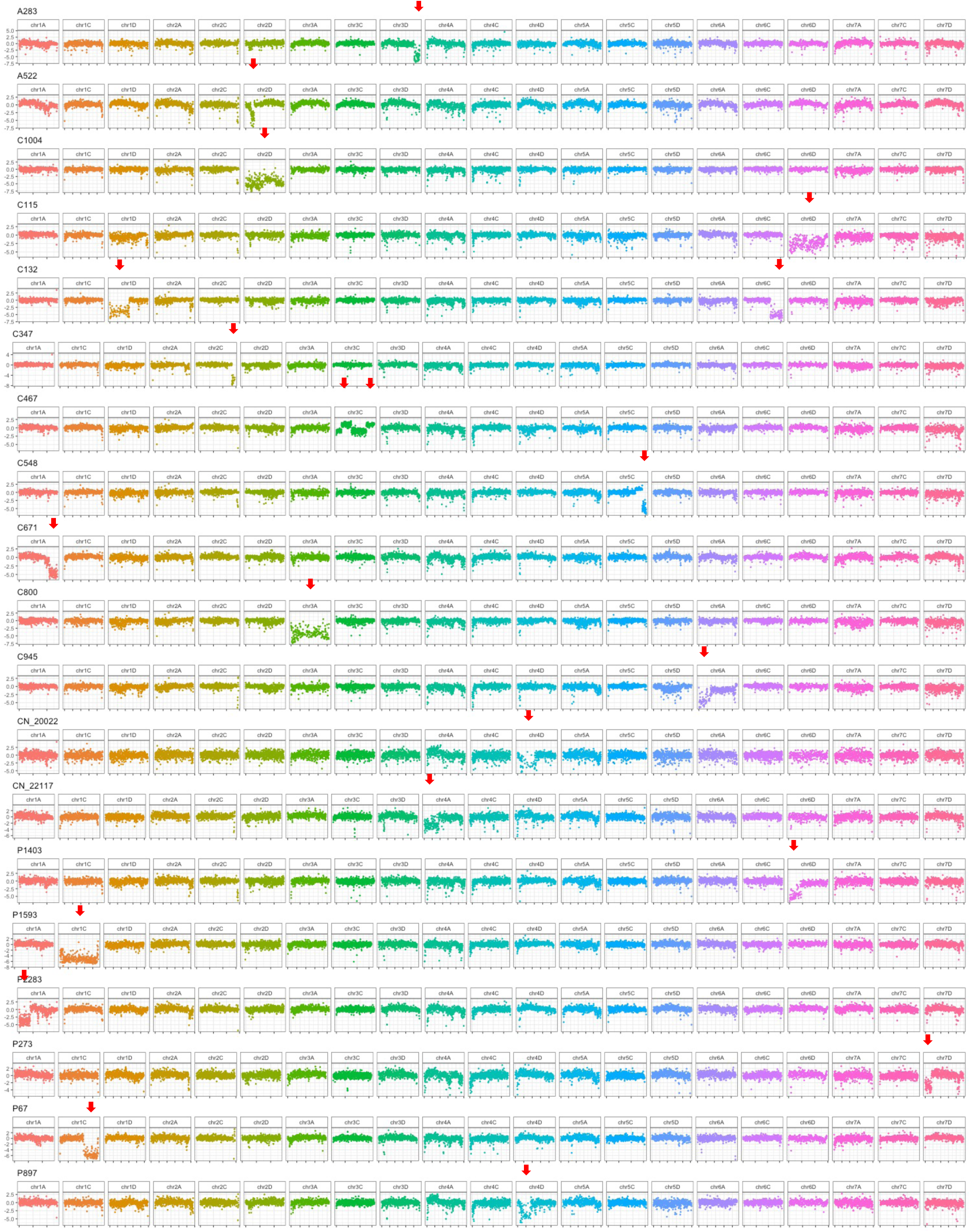
Chromosomal anomalies in representative genotypes from the Global Oat Diversity (G.O.D) panel. G.O.D GBS data were aligned to the GS7 reference genome and normalized to GBS data from GS7, reads were counted in 1 Mb windows. Each row shows one genotype (**Supplementary Table 18**, G.O.D). Red arrows mark SVs. At least one chromosome in each genotype is affected by large SVs, which are most likely deletions, duplications or homeologous exchanges as in the example elaborated on in Fig. 3. A283 is an A. sterilis accessions from Morocco. The read depth variant on 3D is shared with A284, also an A. sterilis accession from Morocco.

**Extended Data Figure 10:**
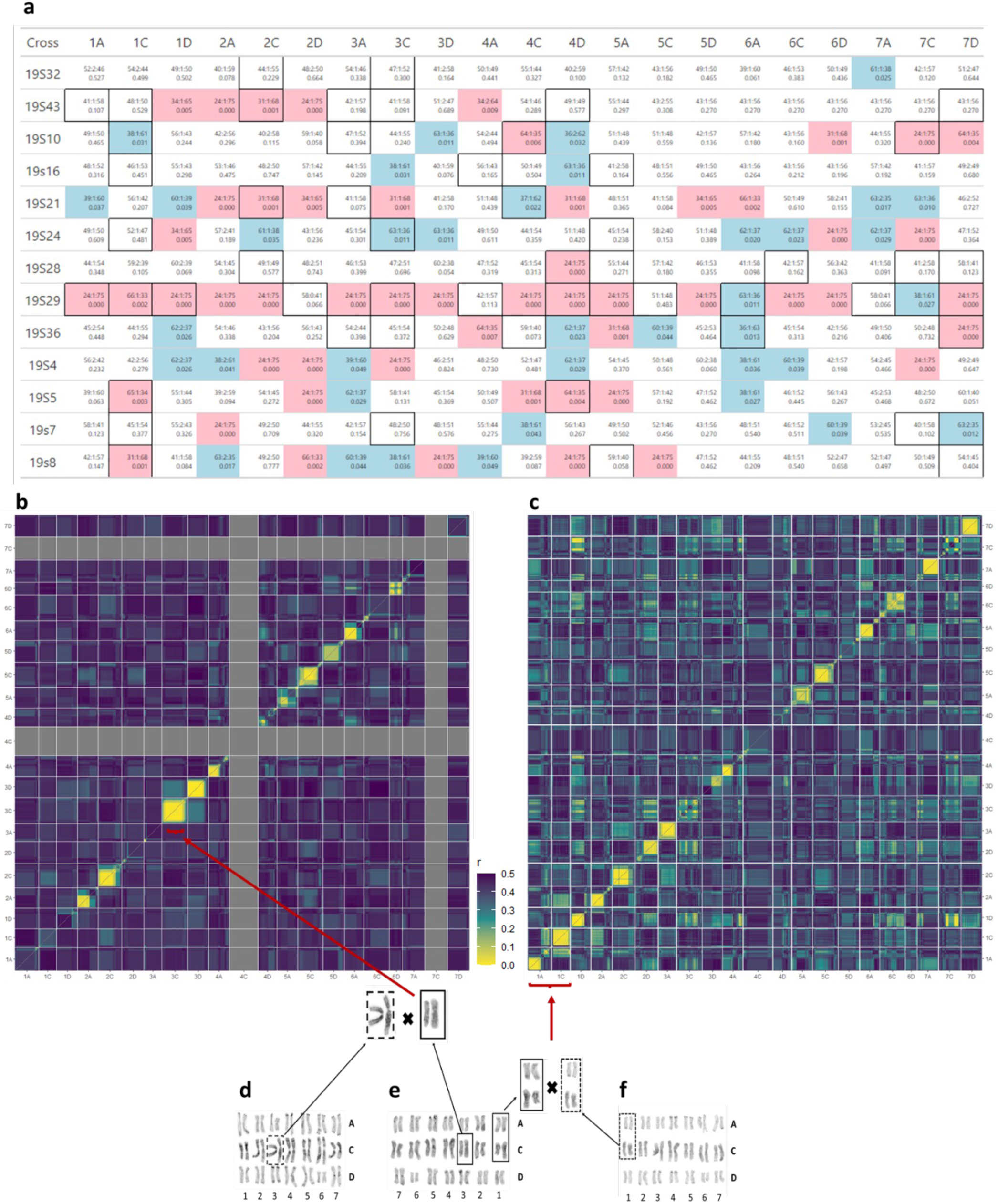
Large-scale chromosomal rearrangements shaped the segregation and recombination patterns in progenies of 13 crosses from a working oat breeding program. **(a)** Summary of 13 crosses (rows) by chromosome (columns). The top row in each cell shows the segregation ratios (AA:AB:BB). The bottom row shows the p-value for the chi-square goodness-of-fit tests of the expected F6 ratio (31:2:31). Cell colors highlight significant P values to reject the fit (pink=p<0.01; blue=p<0.05). Boxes around cells represent suppressed recombination within or between chromosomes. Most crosses demonstrated expected segregation ratios and recombination patterns across the majority of chromosomes. Cross 19S29 showed segregation distortions across most chromosomes. All crosses had at least one chromosome that showed distorted segregation. The recombination patterns of half-sib crosses 19S32 and 19S43, which share the common parent OA1613-5, were investigated in further detail using recombination heatmaps. **(b)** The 19S32 progenies (n=147) exhibited some suppressed recombination, but only on chromosomes 2C and 3C. There were insufficient markers to generate accurate scaled r values on chromosomes 4C and 7C. **(c)** In contrast, the 19S43 progenies (n=122) exhibited pseudo linkage between chromosomes 1A and 1C, as well as between chromosomes 1D and 7C. Additional chromosomes that exhibited some recombination suppression include chromosomes 2C, 3A, 3C, and 4D. Karyotypes of the three parental lines confirmed that they have a complete set of 21 chromosome pairs. **(d)** The karyotype of OA1623-2, the female parent of 19S32, confirmed the presence of a heterozygous inversion on chromosome 3C as well as a homozygous chromosome 1A/1C translocation. **(e)** The karyotype of OA1613-5, the pollen donor for the two crosses, shows a homozygous 3C inversion (non-ancestral) and a homozygous 1A/C translocation. This is manifested in cross 19S32 by a partially suppressed recombination pattern on chromosome 3C and by the expected recombination patterns in 1A and 1C. **(f)** The karyotype of OA1568-6, the female parent of 19S43, shows a pair of ancestral 1A chromosomes (without the 1A/1C translocation), confirming the reason for pseudo-linkage between chromosomes 1A and 1C in cross 19S43. These crosses were not made intentionally for this purpose; rather, they are part of a breeding program that uses both conventional and genomic selection. Even though these populations were small and were advanced by modified single-seed descent, they still show evidence of known SVs and suggest additional SVs in oats. The companion study (Bekele et al., 2024), which detected chromosome inversions using a population-based approach, identified large-scale chromosomal inversions in all affected chromosomes except 7C. A pseudo-linkages similar to 1A/1C between 1D and 7C in 19S43 **(c)** suggests the possibility of an additional translocations.

## Online Methods

### DNA extraction

#### Plant growth and high molecular weight DNA isolation

High molecular weight (HMW) DNA was extracted from young leaf tissue from a single two-week-old seedling grown in an isolated growth chamber under a 12-h photoperiod. The growing temperatures ranged from 18°C (night) to 20°C (day). The hydroponic growth solution was prepared using MaxiBloom® Hydroponics Plant Food (General Hydroponics, Sevastopol, CA, United States) at a concentration of 1.7 g/L. In preparation for PacBio HiFi sequencing, high molecular weight (HMW) DNA was extracted from 72-h dark-treated leaf samples using a CTAB-Qiagen Genomic-tip protocol as described by Vaillancourt and Buell (2019)^1^. DNA quantification and purity were checked using Qubit dsDNA HS assay and Nanodrop spectrophotometer, respectively.

#### Short-read sequencing

Six assemblies included in PanOat have already been published *A. insularis* - BYU209, *A. longiglumis* - CN58138 and Sang^2^; *A atlantica* and *A. eriantha*^3^; and OT3098 which was made available as a free resource by PepsiCo in 2020 and was later improved to pseudomolecules (https://wheat.pw.usda.gov/GG3/graingenes-downloads/pepsico-oat-ot3098-v2-files-2021, PRJEB76239, PRJEB46951).

The *A. byzantina* PI258586 contig assembly was assembled using the TRITEX^4^ pipeline and Hi-C data from Dovetail Omni-C platform. Genome assemblies for Gehl and AC Nicolas were scaffolded using TRITEX and Hi-C to guide pseudomolecule assembly^4^.

### PacBio HiFi sequencing

#### DNA library preparation and Pacbio HiFi sequencing

High molecular weight genomic DNA was sheared to 17 kb on a Diagenode Megaruptor and then converted into SMRTbell adapted libraries using SMRTbell Express Template Prep Kit 2.0. Size selection was performed using a Sage Blue Pippin to select fragments greater than 10kb. These were then sequenced at the BYU DNA Sequencing Center (Provo, UT, USA), except for the ‘Aslak’ accession, which was sequenced at the DNA sequencing and genomics laboratory, (Institute of Biotechnology, University of Helsinki, Helsinki, Finland), using Sequel II Sequencing Kit 2.0 with Sequencing Primer v5 and Sequel Binding kit 2.2. Run times were for 30-hours with adaptive loading, following PacBio SMRT Link recommendations.

#### Hi-C sequencing

*In situ* Hi-C libraries were prepared from young seedlings according to the previously published protocol, using *Dpn*II for the digestion of crosslinked chromatin^2^ or with Phase Genomics multi enzyme mix. Sequencing and Hi-C raw data processing were performed as previously described^5^.

### Genome sequence assembly and validation

#### Chromosome-scale assembly TRITEX + DovetailOmni-C

Chromosome-scale sequence assembly proceeded in three steps: (i) scaffold assembly using the TRITEX pipeline^6^; (ii) super-scaffolding with the Dovetail HiRise pipeline^7^ using Omni-C data; (iii) arranging super-scaffolds into chromosomal pseudomolecules using TRITEX (https://tritexassembly.bitbucket.io). PE450 reads were merged with BBmerge^8^, error-corrected with BFC^9^, and assembled with Minia3^10^. Scaffolding and gap filling were done with SOAPDenovo2^11^ using MP6 and MP9 data. Super-scaffolds were generated with the Dovetail HiRise pipeline from alignments of Omni-C data to scaffolds. Omni-C reads were aligned to the HiRise super-scaffolds with Minimap2^12^. Alignment records were converted to binary Sequence Alignment/Map format using SAMtools^13^ and sorted with Novosort (http://www.novocraft.com/products/novosort/). A list of Omni-C links was extracted from Hi-C alignments using TRITEX scripts. Omni-C links and guide map alignments were imported to the R statistical environment^14^ and analyzed further using TRITEX scripts. An initial Hi-C map was generated using the minimum spanning tree algorithm described by Beier *et al.* (2017)^15^. The assembly and Hi-C map were iteratively corrected by inspecting Hi-C contact matrices, guide map alignments and physical coverage Hi-C reads. Sequence files in FASTA format and AGP tables for pseudomolecules were compiled using TRITEX scripts. The pseudomolecules of *A. byzantina* were aligned against the pseudomolecules constructed from a long-read sequence assembly of cv. OT3098. The OT3098 pseudomolecules (version 2)^16^ were downloaded from GrainGenes^17^.

#### PacBio HiFi

PacBio HiFi reads were assembled using hifiasm (v0.14.1)^18^ and the TRITEX pipeline^6^ was used for pseudomolecule construction. Chimeric contigs and orientation errors were identified through manual inspection of Hi-C contact matrices^4^. GMI423 was used as the reference to map HiFi contigs, using a reduced single copy genome.

*note on chromosome 7D: While assembling chromosome 7D we noticed that, when aligning several of our genotypes to GMI423, there was a large ∼450Mb inversion and a small ∼50Mb sequence with the same orientation as GMI423. We decided to flip the long sequence to the same orientation as GMI423 and flip the small sequence to the inverted orientation, thinking that the small segment was translocated from one end of the chromosome to the other. Retrospectively, this was a mistake. The more plausible explanation would be an inversion of the large sequence, as supported by several genetic studies^19^ showing a distinct lack of recombination in this region.

#### Principal component analysis of GOD GBS data

PCA analysis was done for 9291 genotypes (including the PanOat accessions) using PLINK^20^ (www.cog-genomics.org/plink/1.9/) with –maf=0.05 and a maximum of 70% missing data. Results were plotted in ‘R’ using the ‘ggplot2’ package.

### Single-copy pangenome

A single-copy pangenome was constructed as described by Jayakodi *et al.*^21^ (https://bitbucket.org/ipk_dg_public/barley_pangenome/) with one modification. MMSeq2^22^ was used with the option “ --cluster-mode ” instead of BLAST for all-versus-all alignment. A minimum sequence identity of 95% was required to accept matches. To estimate the pan-genome size the lengths of the largest sequences in each cluster were summed up.

### PanOat transcriptome sequencing

#### Plant materials, growing conditions and tissue dissection

A subset of 23 PanOat genotypes was selected for transcriptome sequencing (**Supplementary Table 2**). RNA was extracted from six tissues (**Extended Data Fig. 2a, b, c** and Fig. 2a). The 23 genotypes were grown in six sets for sampling each tissue separately. Each set comprised at least nine biological replicates (different plants) per oat genotype. Every set with replicates was grown in a separate unit of the growth facility and allocated randomly using the ‘sample’ function in the R statistical environment^23^. Sampled tissues from three different plants (three technical replicates) were pooled into one tube, making one biological replicate and this process was repeated two more times to collect a total of three biological replicates for each of the 23 PanOat genotypes chosen and the six selected tissues (**Extended Data Fig. 2a, b, c)**.

1. **Embryonic tissues**: Seeds were sterilized in ethanol (70%) and sodium hypochlorite (50%) then rinsed five times in sterile water, followed by germination of dehulled seeds in Petri dishes (50mm; covered in two layers of aluminium foil to maintain darkness) in a growth chamber under constant temperature (about 18°C), humidity (about 75% relative humidity), and 16 hour days. Parts of the coleoptile, mesocotyl and seminal roots were dissected from germinating seeds starting from four days after germination (**Extended Data Fig. 2b**). These were promptly frozen in liquid nitrogen and stored at −80°C before thawing prior to RNA extraction.
2. **Leaf tissue**: Seedlings were germinated from sterilized seeds as above, but larger Petri dishes (120 mm) were used. Seedlings were grown until two leaves had emerged. Then, the middle part of the leaf blade was dissected for RNA extraction.
3. **Root tissue**: Seedlings were grown in a small pot on a perlite substrate until three leaves had emerged (**Extended Data Fig. 2b**). Roots were then separated from the perlite and rinsed in sterile water. Cleaned roots were dissected from the top parts of the plants and stored at −80°C until RNA extraction.
4. **Stem tissue**: Plants were grown in pots (1 seed per pot) on a soil substrate in a greenhouse chamber with constant temperature (20°C) and semi-controlled light conditions (16 hours light period) until the main stem and four tillers had developed (**Extended Data Fig. 2b**). Two millimetre-wide stem discs were dissected from the internode elongating below the flag leaf.

5. **Panicle tissue**: Plants were grown as above until the main stem and five tillers had developed (**Extended Data Fig. 2b**). A developing panicle with a size not longer than 15 mm was dissected from the main tiller.

6. **Caryopsis tissue**: Plants were grown as above until the phenophase in between kernel water ripe with no starch and early milk (**Extended Data Fig. 2a, b**). This phenophase is recognized to happen four days post anthesis, according to Ekman *et al*., (2008)^24^.

#### RNA extraction

Total RNA from embryo tissues, leaves, roots, stem and developing panicle was extracted using RNeasy Plant Mini Kit (Qiagen). Total RNA from developing caryopsis tissues was extracted using the RNeasy PowerPlant Kit (Qiagen), according to the manufacturer’s instructions. Prior to RNA extraction, all samples were digested using RNase-free DNase (Qiagen). Tissue samples were thawed and processed in random order. Extracted RNA was diluted in 100 ul of buffer and checked for degradation, quantity and purity. RNA integrity was checked using an Agilent Bioanalyzer. Purity (absence of contaminating proteins) was checked by measuring the fluorescence absorbance of nucleic acids at 260 and 280 nm using a NanoDrop spectrophotometer. RNA amounts were determined using a Qubit fluorometer (Thermo Fisher). Average RNA integrity numbers (RIN) varied from 7.62 in leaf tissues to 9.50 in developing panicles and stem tissues. RIN was on average, lower in leaf tissues but varied little between samples. Only pure RNA samples with high RIN scores (> 8.5; except leaves) and sufficient concentration were used for further processing.

#### Illumina RNAseq

Sequencing libraries were prepared for 432 high quality total RNA samples (RIN >7.62). First, 500-1000 ng of total RNA were Poly(A+) enriched, then RNAseq libraries were produced using the Corall mRNASeq V1 kit according to the manufacturer’s instructions (Lexogen, Vienna, Austria) For each library, barcoding was done by utilizing unique dual indices (UDI). To avoid any experimenter’s bias, the preparation of the libraries was conducted randomly. Sequencing was done in eight pools, with each pool containing 54 randomized single libraries in equimolar amounts. Before sending the pools to the sequencing facility, each pool was sequenced on the iSeq 100 benchtop sequencer at LUKE, Jokioinen, Finland, for quality control. Paired-end sequencing (2x150 bp) was done on a Novaseq 6000 device (Illumina, San Diego CA, USA) distributed on two full S4 flow cells at the Finnish Functional Genomics Centre in Turku, Finland. Sequencing (2x150 bp) of nine libraries was repeated on a NextSeq 550 device (Illumina, San Diego, CA, USA) in the genomics laboratory in Jokioinen, LUKE, Finland. The total number of raw reads per sample and the BioSample IDs are provided in Supplementary **Table 20**.

#### PacBio IsoSeq

For each genotype total RNAs from all tissues and replicates from the respective genotypes were pooled, with between 1623 ng and 2001 ng of pooled RNA used for each library. In total, 24 full length cDNAs were prepared using TeloPrime Full-Length cDNA kit (Lexogen, Vienna, Austria). Different from the manufacturer’s protocol was the purification of 100 µL of cDNA was done with 86 µL ProNex beads (Promega, Madiso, WI, USA), the standard size selection was done according to the Iso-SeqTM Express Template preparation protocol (PacBio, San Diego, CA, USA), and no enrichment for shorter or longer transcripts was utilized. Due to the 5’ cap specificity of this method, only full length, double stranded cDNA was obtained. The cDNAs ranged in size from 1000 to 5800 bp with mean peak values between 1845 bp in Hifi and 2832 bp in GMI423. Following purification, the cDNAs were quantified with Qubit (Thermo Fisher). According to the Iso-SeqTM Express Template preparation protocol (PacBio, San Diego, CA, USA) the amount of cDNA should be in the range of 160-500 ng for Sequel II systems. Libraries with a lower amount of cDNA were re-amplified following the PacBio guidelines. After DNA damage repair, end repair /A-tailing, overhang adapter ligation, and clean up, the concentrations were checked using Qubit. The quality was verified using a Bioanalyzer (Agilent). Twenty-four cDNA SMRTbell libraries with a mean fragment length distribution between 2155 bp and 3557 bp were transferred to the sequencing facility. The Iso-Seq SMRTbell libraries were sequenced at BYU, USA, each library in a separate sequel II run. Numbers of reads and total read lengths are provided in **Supplementary Table 1**.

### Annotation of Protein Coding Genes

For the 23 oat lines with native transcriptome data generated in this study (**Supplementary Table 2**), we performed *de novo* structural gene prediction, confidence classification, and functional annotation, following the protocol described by Mascher, *et al.* 2021^25^. The strategy applied in this study only differs in the use of TE soft-masked genome sequences instead of TE hints (see “Repeatmasking for Gene Detection”). We applied the same gene prediction procedure for the *de novo* annotation of the lines A. *sterilis* TN4 and *A. byzantina* PI 258586 using transcriptome data as evidence. Gene predictions for the lines OT380, *A. sterilis* - TN1, *A. fatua* - CN 25955, *A. eriantha* - BYU132/CN 19328 and *A. atlantica* - Cc7277 which had no native transcriptome data were carried out using a gene consolidation approach described by White *et al.* 2024^26^. Here, the gene predictions for all 30 oat lines described above were cross-mapped with the genome sequences of each other to identify and correct for missed gene models and to annotate genomes without native transcriptome data.

Finally, for the three lines Leggett, Williams and AC Morgan, we predicted their gene content using the projections of the aforementioned evidence-based gene models to their genomic sequences. The principle of the projection method is described in https://github.com/GeorgHaberer/gene_projection; applied parameters of the workflow and code have been deposited in the directory *panoat* of the parent directory.

### Repeatmasking for Gene Detection

To minimize the inclusion of transposon-related gene models, the genome assemblies were softmasked for transposons (TEs) before gene detection. The TE-library used, developed for the oat reference genome^2^, masked approximately 60% of the assembly for each line. Softmasking was performed using vmatch (anaconda.org/bioconda/vmatch, v2.3.0) with the following parameters: “-l 75 -identity 70 -seedlength 12 -exdrop 5 -d -p -qmaskmatch tolower”.

### Construction of the oat core-/shell- and cloud-genome

Phylogenetic Hierarchical Orthogroups (HOGs) based on the primary protein sequences from 30 oat lines with consolidated gene predictions were calculated using Orthofinder version 2.5.5^27^ with standard parameters (see section “Annotation of protein-coding genes” for details; ‘Leggett’, ‘Williams’ and ‘AC Morgan’ were not part of this orthologous framework, as their gene content was not consolidated). Before the analysis, input sequences were filtered for transposon- and plastid-related proteins and proteins encoded on unanchored contigs were discarded for this analysis.

Depending on the focus of the analyses we either treated each of the subgenomes of hexaploid and tetraploid oat lines either as individual entities or, for our analysis of core-, shell- and cloud-genome, as parts of the single lines.

The scripts for calculating core-, shell-, and cloud-genes are deposited in the repository https://github.com/PGSB-HMGU/BPGv2.

Core HOGs contain at least one gene model from all compared 30 oat lines. Shell HOGs contain gene models from at least two oat lines and at most 29 oat lines. Genes not included in any HOG (“singletons”), or clustered with genes only from the same line, were defined as cloud genes.

GENESPACE^28^ was used to determine syntenic relationships between the chromosomes of all 30 oat lines.

### Protein functional annotation and gene-set enrichment analysis

For functional enrichment analysis in the identified expression level categories, Mercator4^29^ (v6.0) protein functional annotation was performed for the identified 5,291 single-copy HOGs across the 20 *A. sativa* lines, which yielded 4,682 protein annotations (**Supplementary Table 21)**. These annotations were used to test enrichment using over-representation analysis (ORA) of sets of genes associated with expression level categories with the R package clusterProfiler^30^ (v4.6) and a Benjamini-Hochberg FDR correction p-value cut-off of 0.05.

Similarly, for ORA across genes classified into core, shell and cloud categories, all the 31 oat lines’ proteomes (2,869,876 proteins and 131,729 HOGs) were functionally annotated with Mercator4 (v6.0). This resulted in a total of 53,018 annotated HOGs for the core (8,325 annotated HOGs), shell (32,108 annotated HOGs) and cloud (12,585 annotated HOGs) categories, applied as universe in the enrichment analyses. ORAs showed enrichment across multiple Mercator4 hierarchical categories (labeled as levels 1-7) https://hmgubox2.helmholtz-muenchen.de/index.php/s/Y3wWa7bn2rayEqw.

### Gene expression analyses

For the analysis of gene expression, RNA sequencing data from 23 oat varieties (**Supplementary Table 2**) were processed using Fastp^31^ for trimming, followed by quality assessment and outlier detection. The data for each line was aligned to its reference genome using Kallisto^32^ and normalized to Transcripts Per Million (TPMs) via Deseq2’s tximport^33,34^ function. All RNA sequencing data were also aligned to the GS7 reference genome for specific analyses.

To compare the expression levels across different subgenomes (A, C, D), gene expression data from six different tissues (leaf, embryo, root, internode, panicle, and caryopsis) were examined. We focused on high-confidence (HC) genes and for each gene, the expression value was normalized using a log transformation (log(value + 1)) to stabilize variance and ensure that the data were suitable for statistical comparisons. The log-transformed expression values were aggregated by calculating the mean expression level for each line, tissue, and subgenome combination. To determine whether the expression levels between different subgenomes were significantly different, Mann-Whitney U tests were performed for each tissue type. Comparisons were made between each pair of subgenomes (A vs. C, A vs. D, and C vs. D).

For the purpose of identifying genes with stable versus variable expression, the analysis was limited to 20 *A. sativa* varieties and 5,965 Hierarchical Orthogroups (HOGs). These HOGs were characterized as single-copy orthologs with an A:C:D ratio of 1:1:1 across all 20 varieties, providing a standardized basis for comparison. HOGs were deemed stable if 90% of the varieties exhibited the same expression category; otherwise, they were classified as variable.

In the set of 5,965 “60-lets”, expression levels were categorized into one of seven categories based on the Euclidean distance to seven ideal expression level profiles: A-, C-, or D-dominant/suppressed, where one gene is predominantly expressed or suppressed, and a balanced category, where A, C, and D genes are equally expressed as outlined in Kamal *et al.* 2022^2^.

### Analysis of gene expression difference between diads and triads

We focused on the 20 *A. sativa* lines and selected genes that had either one single-copy homeoelog in each of the subgebomes (so forming complete triads: A:C:D, 1:1:1 constitution) or had a constitution where one triad member was missing (so forming diads: A:C:D 1:0:1, 1:1:0, or 0:1:1). The lines were then categorized into groups with complete triads and diads, while ensuring uniformity in the missing pattern across the lines. This approach allowed for a controlled comparison across different genetic backgrounds. To ensure robust statistical analysis, specific filtering criteria were applied. Each group analyzed was required to consist of at least five lines, allowing us to achieve sufficient statistical power.

We employed an unpaired t-test to assess the significance of expression differences between groups with a missing homoeolog and those with complete triads. Furthermore, Cohen’s D was utilized to determine the directionality of these differences. A Chi-squared test was conducted to compare the frequency of significant compensatory expressions across the different subgenomes.

### Analysis of the cellulose synthase and cellulose synthase-like gene families

Using previously identified genes encoding for the cellulose synthase superfamily^2^, members of the cellulose synthase and cellulose synthase-like gene families were identified in the 23 oat lines for which RNAseq data was available (**Supplementary Table 2**) using a combined approach of BLAST and orthology analyses using Orthofinder^27^.

To examine whether the deviation in gene copy number contributed directly to variation in expression, the expression of all genes of each subfamily and lines was summed and high and low expression outliers were identified.

Moreover, we sought to identify the major contributors to gene expression within each subfamily and line. Therefore, the percentage expression data for each gene within a subfamily and line combination was identified. This data represented the contribution of individual genes to the overall expression of their respective subfamilies. To detect major contributors, the Elbow method was applied^35^. This method involved sorting the genes by their percentage contribution in descending order and then identifying the point where the largest drop in contribution occurs (the “elbow” point). Genes contributing above this elbow point were considered major contributors, while those below were considered minor contributors. The cumulative percentage contribution for both major and minor contributors was calculated, and a statistical comparison between major and minor contributors using the Mann-Whitney U test was performed to assess whether the difference in contributions was statistically significant.

To assess differences in gene expression between cultivars and non-cultivars across the cellulose synthase and cellulose synthase-like gene families, the mean expression for each gene family within these two groups was calculated. Gene expression data, standardized as z-scores, was calculated for each line. To determine if there were statistically significant differences in gene expression between the two groups (cultivars vs. non-cultivars) within each subfamily, the Mann-Whitney U test was employed. This non-parametric test was chosen due to its ability to compare differences between two independent groups without assuming a normal distribution of the data. For each subfamily, the test produced a U-statistic and a p-value, indicating the likelihood that the observed differences occurred by chance. Given the multiple subfamilies analyzed, the p-values were corrected for multiple comparisons using the Benjamini-Hochberg procedure.

### Whole genome sequencing of North American Spring Oat collection

#### CORE samples sequencing and mapping and GWAS

295 North American spring oat accessions from the CORE population^36^ (**Supplementary Table 15**) were sequenced using an Illumina Novaseq 6000 (paired-end 150bp) with mean read depth of 4.58 per accession.

The genome assembly of GS7 was chosen as a reference as it is a North American variety with a long-read assembly.Adapter sequence (‘AGATCGGAAGAGC’) was removed using cutadapt^37^ and then all reads were aligned to the GS7 reference genome using minimap2^12^, sorted with novosort (https://www.novocraft.com/products/novosort/) and converted to a Compressed Reference-oriented Alignment Map (CRAM^38^) file using SAMtools^39^. A VCF file was created using bcftools ‘mpileup’^39^, included all variations with mapping quality higher than 40. Read depth variation was determined by counting how many reads were aligned in 1 Mb windows along the genome.

For GWAS, a kmer-based reference-free pipeline was used (kmerGWAS^40^). The phenotypes used included heading date, plant height and grain yield, collected from ten locations in two consecutive years (2010 and 2011^36^). K-mers with significant association (-log10 threshold for 10% family-wise error rate) were mapped to the GS7 genome.

#### PanOat assemblies

To align the PanOat assemblies to the GS7 genome, we simulated short reads (10-fold coverage) using fastq generator (https://github.com/johanzi/fastq_generator) and mapped these to GS7 using minimap2^12^. The resulting mapping files were merged into a VCF file together with all 295 CORE genotypes using bcftools merge^39^.

#### Genome alignments

Whole genome alignments of complete pseudomolecule assemblies were performed using minimap2^41^ with the -f 0.05 option to filter out repetitive minimizers and speed the alignment process. Visualization was done using NGenomeSyn^42^ as shown in Fig. 1c.

#### Principal component analysis of WGS data

Focusing only on SNPs on chromosome 7D, PCA analysis was conducted with all 295 genotypes and PanOat assemblies using PLINK with –maf=0.05 and a maximum of 70% missing data. SNP haplotypes were analysed using a custom Perl script^43^ (https://github.com/guoyu-meng/barley-haplotype-script/tree/main/05.other/SNP_haplotype_plot), then sorted by the predicted inversion state from the PCA analysis and plotted in ‘R’.

### Reciprocal translocation 2A/2C in Australian and Canadian Oat

#### Plant materials

The oat varieties used in this study were part of a worldwide oat germplasm collection from the Western Crop Genetics Alliance at Murdoch University (**Supplementary Table 16**), and included 564 lines from 41 countries. To track down the pedigree of the chromosome 2A/2C translocation, potential parental lines for the varieties with the translocation were collected for the second-round test with 32 varieties (**Supplementary Table 17.2**). Seeds of the oats were grown in pots in a glasshouse, with natural lighting cycles and regular watering. Leaves from three-week-old seedlings were collected for DNA extraction.

A recombinant inbred line population was derived from Bannister / Williams crossing. An F_5_ population was grown in the greenhouse at InterGrain Pty Ltd (19 Ambitious link, Bibra Lake, WA 6163). A total of 188 lines, together with their parents, were used for molecular map construction using DarT seq technology, following the online instruction (Diversity Arrays Technology Pty Ltd, Canberra, Australia). Three to five centimetres of leaf sections were collected for DNA extraction from seedlings at the three-week stage.

#### Chromosome specific molecular markers for 2A/2C translocation

Genomic DNA was extracted from the leaves of each oat line using the cetyl-trimethyl-ammonium bromide (CTAB) method^44^. DNA quality was assessed on 1% agarose gels and quantified using a NanoDrop spectrophotometer (Thermo Scientific NanoDrop Products, Wilmington, Delaware USA). DNA was diluted to 50ng/µl for PCR.

Two DNA samples (OT207 and Kanota) used in this study were obtained from Agriculture & Agri-Food Canada, Ottawa, due to the unavailability of these varieties in Australia. Specific primers for PCR (**Supplementary Table 17.1)** were based on sequences from the Australian oat Bannister and the Spanish oat FM13. Primers Ban2A_F3 and Ban2A_R3 are specific to the Bannister chromosome 2A breakpoint, producing a PCR amplicon of 397 bp. Another pair of primers, Ban2C_F3 and Ban2C_R3, are specific to the Bannister chromosome 2C breakpoint, producing a PCR amplicon of 595 bp. PCR was carried out in 10 µl reactions in 384-well PCR plates (Axygen, USA) containing 50 µM of each of the three primers, 200 µM dNTPs, 1.5 mM MgCl2, and BIO-TAQ (Biolne Australia) in a Veriti Thermo cycling machine. The PCR products were separated and visualised on 2% agarose gels stained with GelGreen® (Biotium, USA). Oat lines containing the 2A/2C translocation produced bands and were scored as “present”, while normal oat lines did not produce bands and were scored as “absent”.

#### Genetic map construction

The RIL lines were genotyped with DArTseq markers. The genotypes were filtered with the following parameters: call rate above 90%, Polymorphic Information Content above 0.2, and heterozygous frequency less than 0.6. MSTmap^45^ (https://github.com/ucrbioinfo/MSTmap) was used to construct the genetic map^45^. Several rounds of calculations were carried out to correct and impute genotype calls. After the first round of genetic map construction, the markers were sorted based on the genetic map and considering the physical orders. Missing data and noisy markers were corrected if the physical and genetic orders were consistent. The heterozygous regions were fixed first, and then the nearby markers were corrected in subsequent rounds of calculation. The final genetic map was calculated after 3–4 rounds of corrections. The chromosomal 2A/2C translocation-specific molecular markers were manually integrated into the molecular linkage map.

#### Validation of chromosome 2A/2C fragment deletion in the RIL population of Bannister/Williams

The whole genome DarT seq method identified ten RIL with potential chromosome fragment deletions (**Supplementary Table 17.3**), but only three RIL had set seeds. Based on DArT marker sequences, PCR primers for different locations on chromosome 2A or 2C were designed to validate the truncations seen using agarose gel-based methods. The primer sequences and locations (aligned FM13 genome sequences) are listed in **Supplementary Table 17.4**. Each pair of primer sequences or a marker is unique and only has one specific amplicon.

PCR and gel analysis was carried out as described above. With this set of primers, a score of “present”, indicated the DNA sequence at a particular locus was normal. Those scored as “absent” indicated mismatches with the primers, or that a sequence was missing. When a few consecutive loci on the same chromosome were all scored as “absent”, it is most likely that the section of the chromosome has been replaced or is truncated.

Sixteen markers scattered across chromosomes 2A and 2C were tested in three lines (BW041, BW080 and BW123). The three or four plants from each pot were verified to have the same genetic background. BW041 and BW080 are likely truncated on 2C from 412,337Kb. Tip of the Chromosome 2A from BW123 is missing; the truncation point is likely between 2A 26, 716Kb and 344,891Kb. The deleted fragments were further validated by 10x whole genome shotgun sequencing.

#### Yield evaluation

Grain yield data were obtained from the 2017-2022 Australian National Variety Tests. Each year included 19 to 31 trials across Australia, with a total of 158 trials designed in three replications. There were six 2A/2C translocated varieties and eleven non-translocated varieties.

#### QTL mapping

Plant height was measured before grain harvest. The genotypic and quantitative trait data were formatted for use with MapQTL5.0. A permutation test was conducted to calculate the LOD value threshold for positive QTL detection. Internal mapping was first performed to identify the markers with the highest LOD value above 3.2. The markers with the highest LOD values from different QTL were selected for Multiple QTL mapping (MQM) analysis.

### Large-scale chromosomal rearrangements shaped the segregation and recombination patterns in progenies of 13 crosses from a working oat breeding program

Thirteen F_1_ crosses were made in 2019 at the oat breeding program at the Ottawa Research and Development Centre, AAFC (**Supplementary Table 22.1**), among oat lines that were selected for their excellent trait profiles and adaptation to Canadian environments. Progenies were advanced by a modified single-seed descent method to the F_6_ generation. Parents and progeny were genotyped using a targeted genotyping-by-sequencing method^46^. Progenies were filtered to remove those with >90% similarity to a parent and to eliminate progeny that showed >98.5% similarity to another progeny. The position of tag-level haplotype markers on the Sang reference genome^2^ was used to compute recombination fractions (r) between all pairs of markers. Values of r were averaged within a sliding window of 20 Mbp at 10 Mbp increments in two directions of a complete genome matrix, as described by Tinker *et al*. 2022^47^, such that recombination fractions were scaled to physical distance. Recombination matrices across the full genome were displayed as heatmaps, colored from yellow (r=0) to cyan (r=0.2) to purple (r>=0.4) (Also in **Supplementary Table 22.2**). Chromosomes with inadequate marker coverage to estimate recombination were colored grey.

### C-banding

All karyotypes of lines throughout the manuscript were determined by C-banding as described by Mitchell *et al.* (2003)^48^, except that 0.1% colchicine at 20 C for 3-5 h was used to arrest microtubule assembly

### Data download and availability

The data generated by the PanOat Consortium are made freely available and publicly accessible through deposition in public databases. Sequence data were deposited in the European

Nucleotide Archive under project IDs PRJEB56828 (genome assembly raw data), PRJEB57570 (transcriptome sequencing) and PRJEB62778 (WGS resequencing data). Project IDs for individual assemblies and BioSample IDs for individual CORE genotypes are listed in **Supplementary Tables 19 and 15**, respectively. The annotation datasets are available for download from the USDA-ARS GrainGenes database^17^ at https://wheat.pw.usda.gov/GG3/content/panoat-data-download-page. This page also serves as a landing page for access not only to data but also to genome browser tools and BLAST services^49^.

#### Genome browsers

33 genome browsers for each PanOat accession were created in GrainGenes. The links to these genome browsers are available from the ‘Data Download’ landing page mentioned above. They are also available from the main GrainGenes Genome Browser landing page at https://wheat.pw.usda.gov/GG3/genome_browser. Each genome browser contains datasets as tracks, which include pseudomolecule sequences, as well as high-confidence and low-confidence gene models. The gene models have external links to the eFP Browser at the University of Toronto [https://bar.utoronto.ca/eFP-Seq_Browser/].

#### BLAST

BLAST services in GrainGenes include databases that have pseudomolecules from 35 accessions, as well as scaffold sequences from a subset of six accessions: BYU960, *A. byzantina* PI258586, Leggett, AC Morgan, OT3098 v2, and Williams. Note that, when a BLAST query sequence hits a region of a genome assembly that has a genome browser in GrainGenes, a clickable link to the hit region on that genome browser is made available through the JBrowse Connect API^50^, as are other details, such as hit scores, statistics, and sequence alignments.

